# Dynamic regulation of protein homeostasis underlies acquired thermotolerance in Arabidopsis

**DOI:** 10.1101/2023.08.04.552042

**Authors:** Mayur Bajaj, Annapurna Devi Allu, Basuthkar J Rao

## Abstract

Rapid climate change demands the development of heat-resilient plants. Elevated temperatures perturb cellular protein homeostasis, and its timely restoration is crucial for plant survival after stress. Thermopriming, which involves pre-exposure to sublethal heat stress, has emerged as a promising strategy for enhancing heat stress tolerance. However, the impact of thermopriming on protein homeostasis remains unclear. Here, we demonstrate that priming-mediated acquired thermotolerance involves the dynamic regulation of protein maintenance and clearance mechanisms. Priming facilitates the activation of heat shock response (HSR) via HSFA1, and unfolded protein response (UPR). Simultaneously, priming induces the protein clearance pathway, namely autophagy, potentially through the dynamic modulation of autophagy-negative regulators. Contrastingly, unprimed seedlings fail to mount HSR and UPR, resulting in disrupted proteostasis and the accumulation of aggregates, and ultimately fail to survive. While the loss of UPR was found to have a minimal impact on priming-mediated outcomes, the HSR response proved essential, as its absence led to lethality under heat stress. Additionally, the absence of HSR was found to enhance the autophagy response post-stress. Our results highlight the critical role of protein maintenance mechanisms over clearance pathways in ensuring survival. Taken together, our study demonstrates that thermopriming enhances heat stress resilience by temporally coordinating autophagy, HSR and UPR responses to maintain proteostasis.

## Introduction

In nature, plants often encounter multiple abiotic and biotic stressors that affect their cellular homeostasis. High temperatures are one such abiotic stressor that is detrimental to plant growth, productivity and even survival. Heat stress reduces protein, starch, and oil content (Fahad et al., 2017) and can lead to substantial economic losses in agriculture (Long & Ort, 2010; Schmitt et al., 2022). Studies have reported that for every 2°C rise in temperature, there could be a 50% decline in crop productivity (Lobell et al., 2012). As sessile organisms, plants cannot escape adverse conditions and must instead preemptively activate protective genes to survive. Recent studies have shown that pre-exposure to a stress stimulus (referred to as ‘priming’) can dramatically improve plant stress tolerance (Liu et al., 2022; Samantaray et al., 2023).

‘Priming’ refers to the pre-conditioning with non-lethal stress treatment used to induce acquired stress tolerance. The priming treatment allows the plant to robustly respond to a subsequent severe stress, referred to as ‘triggering’ (Bäurle, 2016; Conrath et al., 2007; Sedaghatmehr et al., 2016; Wilkinson et al., 2019). Several stimuli, including mild environmental stress factors, hormones or chemical moieties can serve as priming agents (Nair et al., 2022). Priming-mediated acquired stress tolerance has been previously studied in the model plant system *Arabidopsis thaliana* (hereafter referred to as Arabidopsis) along with several crop plants (Balmer et al., 2018; Liu et al., 2022; Samantaray et al., 2023; Song et al., 2015; Ye et al., 2019). Priming can aid in remodeling the transcriptomic, epigenetic, or metabolic programs, thereby increasing the robustness of plant responses to subsequent stress (Catoni et al., 2022; Espinas et al., 2016; Gamir et al., 2014; Serrano et al., 2019; Tugizimana et al., 2018). Although several studies have focused on understanding the transcriptional and epigenetic regulation underlying the priming response (Crespo-Salvador et al., 2018; Kotkar & Giri, 2020; Nair et al., 2022; Noh et al., 2021), very little is known about the role of post-translational processes such as protein homeostasis, autophagy and mitochondrial dynamics, which are critical for plant survival (Raffeiner et al., 2023; Selinski et al., 2024; Sun et al., 2021).

Heat stress primarily disrupts protein homeostasis by destabilizing thermolabile proteins (Richter et al., 2010). In response, plants activate multiple proteostasis pathways - including molecular chaperones, autophagy, and the ubiquitin-proteasome system (UPS) - to restore protein homeostasis and promote stress tolerance. Upon heat stress, misfolded proteins accumulate thereby activating heat shock response (HSR) and unfolded protein response (UPR) (Gao et al., 2023). HSR involves accumulation of heat shock proteins (HSPs), which facilitates the refolding and disaggregation of misfolded proteins (Richter et al., 2010; Rosenzweig et al., 2019). *HSFA2* and *HSFA3* are well-established HSR molecules involved in transcriptional upregulation of chaperones under acclimatizing heat stress conditions (Friedrich et al., 2021). The UPR in plants is a complex endoplasmic reticulum (ER)-associated pathway which involves activation of multiple transducers, including members of the IRE1, bZIP and NAC families to resolve proteotoxic stress (Nawkar et al., 2018; Pastor-Cantizano et al., 2020). Additionally, proteotoxic stress also activates degradative pathways such as UPS and autophagy. Where UPS is involved in clearance of unfolded/misfolded proteins, autophagy contributes by removing damaged organelles and promotes the recycling of dysfunctional proteins/aggregates during heat stress (Raffeiner et al., 2023; Sharma et al., 2024; Su et al., 2020; Zhou et al., 2013). Autophagy is tightly regulated by signaling pathways involving TOR, SnRK, and IRE1. TOR (target <u>o</u>f <u>r</u>apamycin) inhibits autophagy under nutrient-rich conditions by phosphorylating components of the autophagy initiation complex (Suttangkakul et al., 2011), whereas, SnRK (sucrose non-fermenting 1 (SNF1)-related protein kinase) promotes autophagy during energy deprivation by inhibiting TOR (Soto-Burgos & Bassham, 2017). IRE1 (Inositol-requiring 1), an ER-membrane protein, is proposed to positively regulate autophagy via its mRNA degradation activity (Bao et al., 2018). Additionally, HSFA1a has also been identified to induce autophagy under heat and drought stress by upregulating *ATG* genes (Wang et al., 2015; Xie et al., 2022). Autophagy also plays a role in resetting long-term heat-stress memory (Sedaghatmehr et al., 2019) by degrading specific chaperones HSP21 and HSP90.1, through FtsH6- and NBR1-mediated mechanisms, respectively (Sedaghatmehr et al., 2021; Thirumalaikumar et al., 2020). However, holistically, how protein homeostasis mechanisms contribute towards priming-mediated responses remain unclear.

In this study, we examined the dynamics and regulation of proteostasis pathways to gain insights into the post-translational landscape during priming-mediated acquired thermotolerance. Our findings suggest that short-term thermopriming enhances tolerance to severe heat stress by concurrently activating protein protective responses through the accumulation of chaperones and folding catalysts via HSR and UPR; and facilitates clearance of damaged proteins via dynamic modulation of autophagy. Conversely, when exposed to heat stress, unprimed seedlings fail to activate protein protective responses, experience translational arrest, accumulate protein aggregates, leading to disruption of proteostasis and ultimately fail to survive. Overall, our study offers novel insights into priming-mediated thermotolerance and highlights the critical role of protein homeostasis in maintaining cellular resilience under heat stress.

## Results

### Priming improves survival from severe heat stress

To understand the mechanisms driving short-term acquired thermotolerance, we exposed the five-day-old Arabidopsis wild-type (WT) seedlings to 37°C (for 90 min) and allowed them to briefly recover at optimum conditions (for 90 min) before exposing them to triggering heat stress (45°C for 45 min) (**Figure 1A**). Hereafter, this condition is referred to as ‘primed’. The effect of thermopriming was analysed by comparing the impact of lethal heat stress on primed *vs.* naive (unprimed: directly exposed to triggering heat stress) seedlings. Post heat stress, seedling growth was recorded as a function of time under all the conditions tested (control, primed and unprimed) (**Figure 1B**). Post-exposure to triggering heat stress, primed seedlings displayed a halt in growth; however, the growth resumed quickly, which was found to be significantly higher than the unprimed, quantified as leaf number, root length and fresh weight (**Figure 1, C-E**). By 5 days post stress (DPS), unprimed seedlings display a visibly complete loss of chlorophyll and fail to survive (**Figure 1B**). In summary, heat stress affects growth in both primed and unprimed conditions. However, primed seedlings can mount a more robust response, facilitating the swift resumption of growth after heat stress.

**Figure 1:**
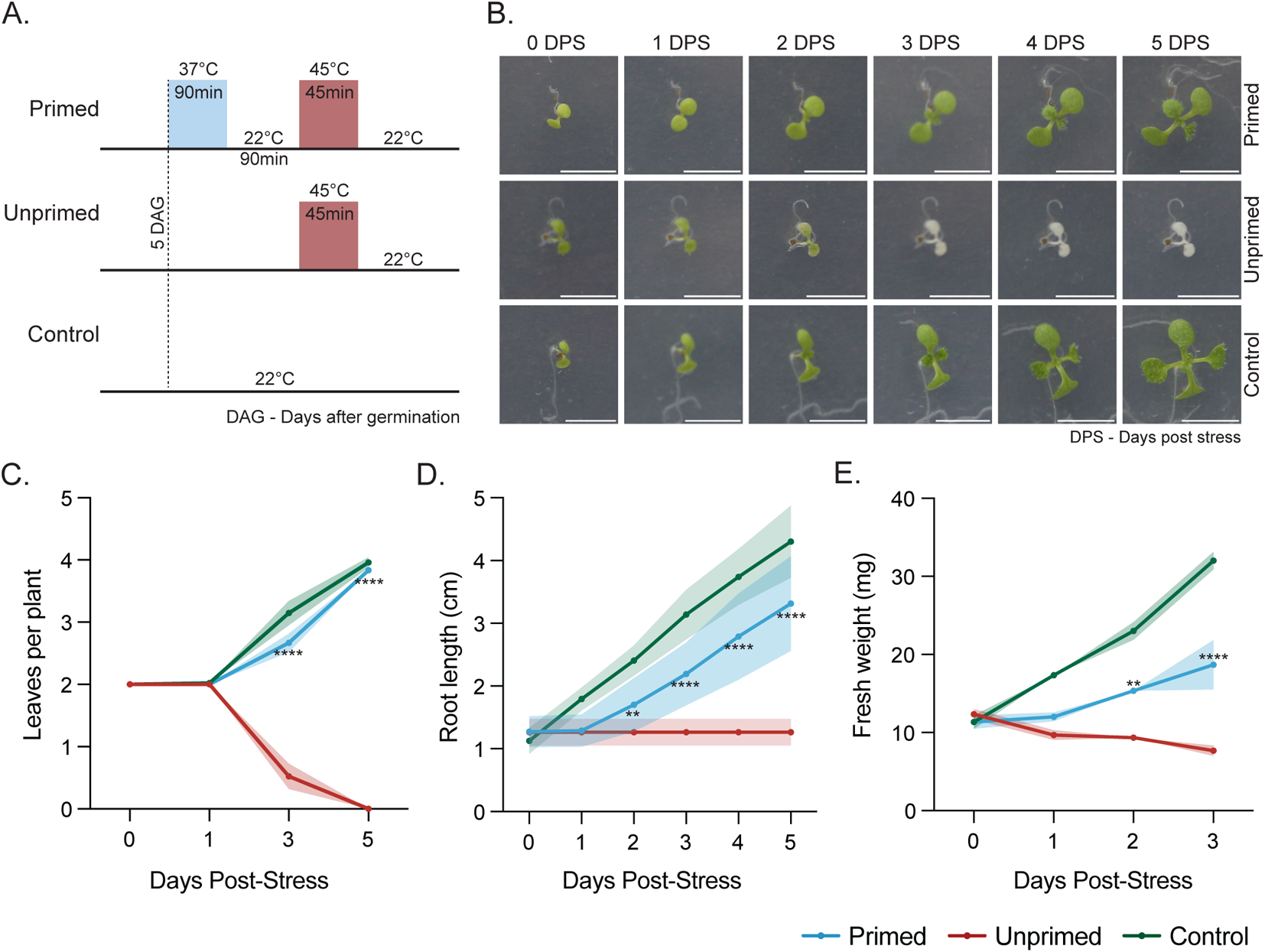
Priming improves survival under severe heat stress. (A) The experimental paradigm is schematised where 37°C was used as a priming stimulus, followed by recovery at 22°C (optimal growth temperature), and then exposed to a 45°C heat-triggering stimulus for the indicated time durations. Control (untreated) and unprimed seed-lings (subjected to only 45°C) were compared in parallel. (B) Post-heat stress, seedlings were monitored over multiple days at 22°C to assess their recovery. The panel depicts representative images for primed, unprimed and control conditions. Scale bar, 0.5cm. The growth recovery was quantified based on (C) the number of leaves per plant (n=3 biological replicates, with 16 seedlings per replicate), (D) root length (n=3 biological replicates, with 6 seedlings per replicate), and (E) fresh weight (n=3 biological replicates, with 16 seedlings pooled per replicate) at the indicated time points. Despite an initial halt in growth (as observed in root length), primed seedlings resumed growth similar to the control, whereas unprimed plants exhibited complete growth arrest and ultimately failed to survive. Data are present-ed as mean ± SEM (for leaves per plant) and mean ± SD (for root length and fresh weight). Statistical significance was determined using two-way ANOVA, where the asterisk (** p <0.01; **** p <0.0001) indicates significant differences between the primed and unprimed seedlings.

### Thermopriming mitigates heat stress-induced protein damage

To determine the impact of heat stress (with or without priming) on protein homeostasis, we analysed the protein content in seedlings across all conditions (**Figure 2A**). The total protein content in the control (grown under optimal conditions at 22°C) and primed seedlings was found to be similar across all the time points analysed post-heat stress (**Figure S1A**). In contrast, unprimed seedlings displayed a consistent decline in total protein levels from 1 to 3 DPS, signifying a reduction in total protein content (**Figure S1A**).

**Figure 2:**
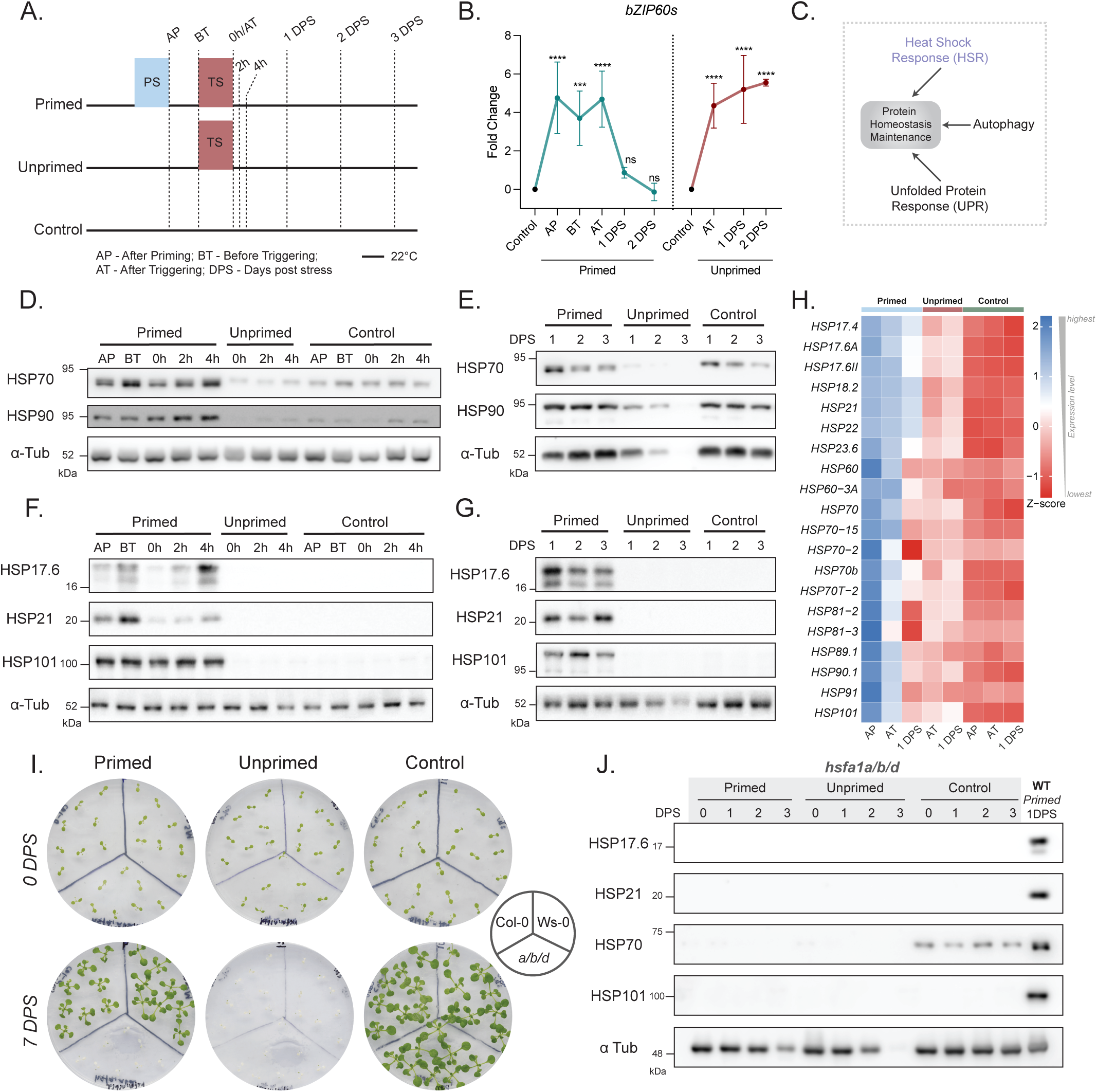
Priming leads to an increase in chaperone abundance post-heat stress via HSFA1. (A) The schemat-ics of the experimental paradigm is shown. AP, BT, and 0-3 DPS denote the time points at which seedlings were collected (PS-Priming Stimulus, TS-Triggering Stimulus). (B) Misfolded protein accumulation following heat stress was determined by checking for the presence of *bZIP60* spliced form (*bZIP60s*). Expression analysis suggested that primed seedlings initially accumulated misfolded proteins but recovered by 1 DPS, whereas unprimed seedlings experienced a sustained accumulation of misfolded proteins (n=3 biological replicates). Statistical significance was calculated using one-way ANOVA (**** p <0.0001; *** p <0.001; ns – not significant). (C) Illustrated representation of the processes involved in the maintenance of protein homeostasis. (D-G) To assess the status of HSR, western blot analysis of HSP17.6, HSP21, HSP70, HSP90, and HSP101 proteins in various samples was performed post-heat stress. The data revealed that priming led to an increase in the abundance of HSPs, which persisted even after stress removal. In contrast, unprimed seedlings did not show any increase in HSP abundance compared with the control (n=3 biological replicates). α-Tubulin served as a loading control. (H) Heat map showing the mean expression levels of *HSPs* across different time points, derived from transcriptome data (n=2 biological replicates). The data suggests that priming stimulus led to an increase in *HSP* levels compared to control. Unprimed seedlings also displayed an increase in *HSP* abundance post-stress compared to control, but the level remained lower than that of primed seedlings. (I) The panel depicts representative images of *hsfa1a/b/d* (referred to as *a/b/d*), Col-0 and Ws-0 post-priming/triggering heat stress. The effect of priming on heat stress response was lost in the *a/b/d* mutant compared to Col-0/Ws-0. Unprimed *a/b/d* failed to survive, similar to that of Col-0/Ws-0 post-heat stress (n=3 biological replicates). (J) Immunoblot analysis of HSPs revealed that primed-WT displayed an increased abundance post-priming, whereas primed *a/b/d* failed to display HSP abundance (n=3 biolog-ical replicates).

Heat stress causes protein denaturation, leading to the accumulation of misfolded/unfolded proteins, which, if not resolved, can lead to severe consequences. Misfolded protein accumulation triggers the splicing of *bZIP60u* (unspliced), forming *bZIP60* spliced form (*bZIP60s*), which is known to transcriptionally modulate genes involved in restoring cellular proteostasis under stress conditions (Deng et al., 2011; Humbert et al., 2012). To assess whether proteotoxic stress was induced in our experimental regime, we analysed the transcript abundance of *bZIP60s* across all the conditions at different time points post-priming/triggering stress (**Figure 2A**). Interestingly, exposure to priming temperature itself triggered misfolded protein accumulation, as evidenced by the presence of *bZIP60s* at AP time point (**Figure 2B; S1, B-C**). Following the triggering heat stress, while both primed and unprimed seedlings accumulated misfolded proteins (elevated *bZIP60s* at AT/0 DPS), only primed seedlings recovered to control levels by 1 DPS (**Figure 2B; S1, B-C**). The unprimed seedlings displayed sustained proteotoxic stress as evidenced by elevated *bZIP60s* levels throughout the recovery phase (**Figure 2B; S1, B-C**). These results suggest that priming-mediated responses may be involved in the efficient maintenance of the protein pool. This prompted us to assess the status of protein homeostasis maintenance pathways (i) heat shock response (HSR), (ii) unfolded protein response (UPR), and (iii) autophagy response during our heat stress paradigm.

### Thermopriming facilitates effective heat shock response mediated by HSFA1

We determined the HSR status (**Figure 2C**), by analysing chaperone levels (HSP70 and HSP90) post-heat stress across all conditions. Exposure to the priming stimulus (AP) resulted in increased accumulation of HSP70 and HSP90, which persists until 1 DPS post-exposure to triggering heat stress in the primed, compared to the control (**Figure 2, D-E**). However, the unprimed seedlings failed to mount such a response, with the levels of HSPs being lower than those of the control seedlings (**Figure 2, D-E**). In addition, we analysed the abundance of other heat stress inducible proteins HSP17.6, HSP21 and HSP101 (Bäurle, 2016; Friedrich et al., 2021) at all the time points stated above. Increased levels of all three HSPs were detected in the primed, but not in the unprimed or control seedlings (**Figure 2, F-G**). Immediately upon exposure to priming (AP), an increased abundance of HSP17.6, HSP21 and HSP101 was detected (**Figure 2F**), which persisted until 3 DPS (**Figure 2G**). Furthermore, to assess whether the HSP abundance observed thus far parallels transcript levels across all conditions, we profiled the expression of *HSPs* using data available from RNA-sequencing (**Figure S2, A-C**), followed by confirmation using qPCR. In the primed seedlings, the abundance of HSPs correlated with their elevated transcript levels compared to those of the control (**Figure 2H; S2D**). Surprisingly, unprimed seedlings displayed a gradual increase in *HSPs* transcript abundance post-heat stress (**Figure 2H; S2D**), which did not correlate with the observed protein levels (**Figure 2, D-G**).

Furthermore, we examined the involvement of HSFA1, a core regulator of the plant heat stress response (Bakery et al., 2024). We profiled the expression of *HSFA1a* and its direct and indirect downstream targets, *HSFA2* and *HSFA3* (Ohama et al., 2017), following heat stress. We observed that *HSFA1a* levels did not change significantly across the time points tested, except at the AT/1DPS in unprimed (**Figure S2E**). These observations are in line with previous reports where *HSFA1a* levels did not change significantly (Löchli et al., 2023), likely because HSFA1’s regulation under heat stress is predominantly controlled at the post-translational level (Mesihovic et al., 2022; Ohama et al., 2017; Wang et al., 2023). In contrast, the downstream targets *HSFA2* and *HSFA3* were upregulated in both primed and unprimed following heat stress (**Figure S2E**). Notably, *HSFA2* induction persisted even at 2 DPS, whereas *HSFA3* induction returned to basal levels by 1 DPS in primed seedlings (**Figure S2E**).

HSFA1s are a well-established HSR regulator involved in transcriptional upregulation of chaperones (Ohama et al., 2017). In our study, we observed HSFA1a’s downstream activity (*HSFA2/HSFA3* levels) to be correlated with the abundance of *HSPs* across our regime. These observations prompted us to assess the impact of HSFA1s loss on the priming-mediated heat stress response. To overcome the functional redundancy of the HSFA1s in Arabidopsis, we used a previously reported *hsfa1a/b/d* triple mutant line (Yoshida et al., 2011) for our analysis. The unprimed *hsfa1a/b/d* and Col-0/Ws-0 did not show any differences in their basal heat stress response (**Figure 2I**). However, even upon priming, the *hsfa1a/b/d* mutants failed to survive following heat stress, unlike the Col-0/Ws-0 seedlings (**Figure 2I**). Furthermore, priming-mediated HSP induction was abolished in the *hsfa1a/b/d* mutant in contrast to the WT (**Figure 2J**). These findings underscore the essential role of HSFA1s in mediating priming-dependent chaperone induction and thermotolerance.

### Unprimed seedlings experience translational arrest following heat stress

Notably, despite the activation of HSFA1 and the concurrent increase in *HSP* transcript levels, unprimed seedlings failed to show a corresponding increase in HSP protein abundance. This prompted us to check the translational status across these conditions. To this end, we examined the expression profiles of differentially expressed genes (DEGs) involved in protein translation using our transcriptome data. We observed that exposure to priming stimulus alone (AP time point) has a minimal impact on the expression levels of genes encoding RNA polymerases and ribosomal proteins. However, exposure to triggering stimulus (AT timepoint) led to a severe downregulation of these genes, indicating a repression of translation (**Figure S3A**). Interestingly, during the heat-stress recovery phase, the expression of these genes was restored to near-control levels in the primed seedlings, whereas, the unprimed seedlings exhibit a sustained downregulated gene expression, suggesting a sustained halt in translation (**Figure S3A**). These findings corroborate our earlier observation that although *HSP* transcript accumulates in the unprimed, the lack of HSP induction is likely due to impaired translation.

### Priming activates the transcriptional unfolded protein response

Dysregulation of protein homeostasis is known to impact ER functional status, triggering the activation of unfolded protein response (Humbert et al., 2012; Nawkar et al., 2018). The UPR aims to restore ER-proteostasis by upregulating genes involved in ER-quality control (ERQC) and ER-associated degradation (ERAD) pathways (Duan et al., 2023; Nawkar et al., 2018). To assess the status of UPR, we examined the expression profiles of DEGs involved in ERQC and ERAD pathways. We observed that priming stimulus alone can lead to an increase in the expression of several canonical UPR components, including members of the bZIP and NAC transcription factor families, as well as luminal binding proteins (BIPs), protein disulfide isomerases (PDILs), ER-resident DnaJ proteins (ERDJs /P58IPK) and others, which are indicative of early ERQC activation (**Figure 3A**). Additionally, the upregulation of various ubiquitin ligases implicated in the ERAD pathway was also observed upon priming (**Figure 3A**). Importantly, this transcriptional upregulation persisted in primed seedlings even at the post-triggering time point. In contrast, despite experiencing heat stress, unprimed seedlings failed to exhibit any increase in the abundance of these transcripts (**Figure 3A**). These results indicate that priming enables transcriptional activation of the UPR in response to heat stress.

**Figure 3:**
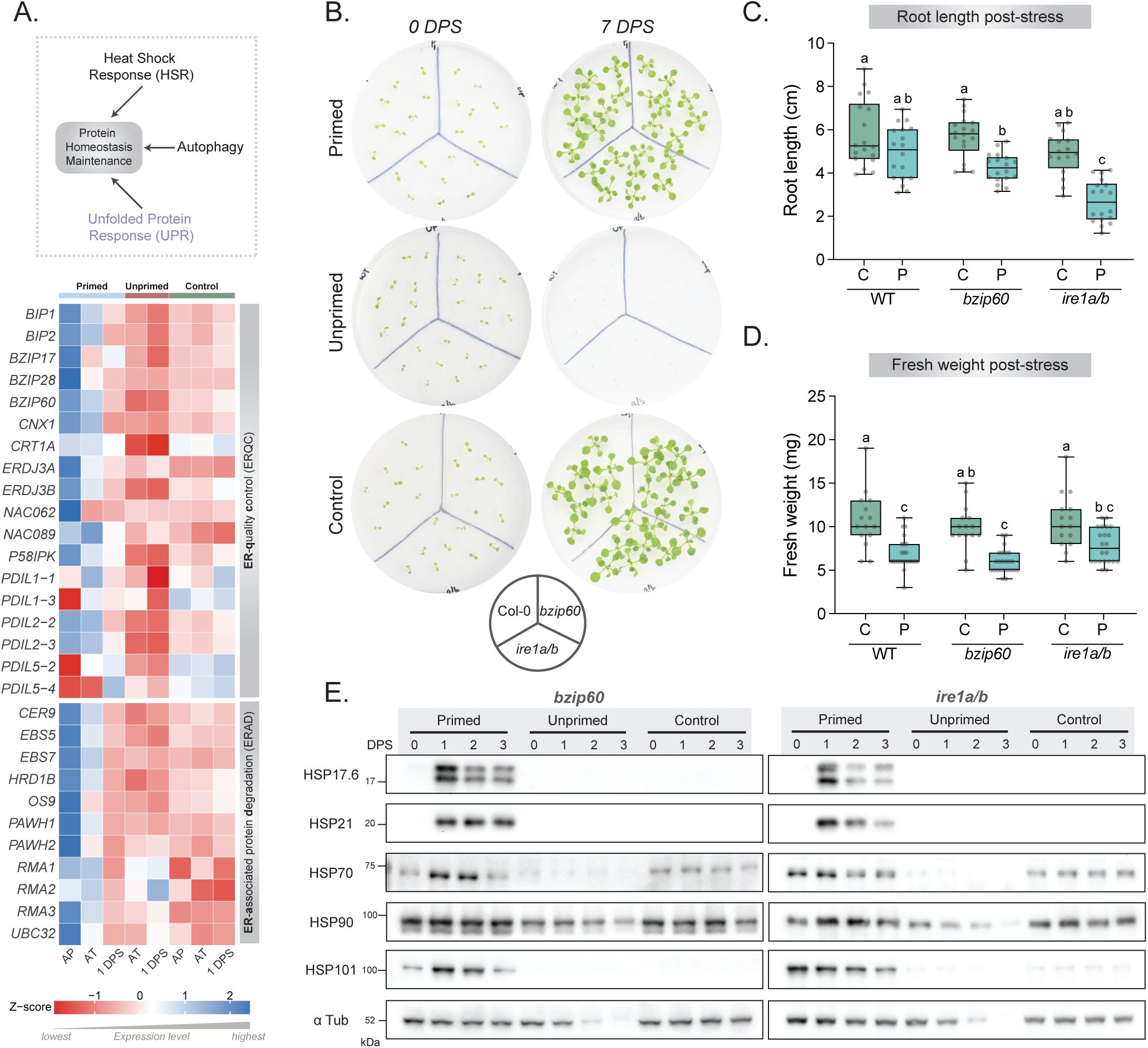
Priming stimulus activates the transcriptional unfolded protein response, while loss of UPR delays seedling recovery post-heat stress. (A) Transcriptome data was used to assess unfolded protein response (UPR) following heat stress. The heatmap displays the mean expression levels of genes involved in endoplasmic reticulum (ER) quality control (ERQC) and ER-associated degradation (ERAD), representing two major branches of the UPR pathway. Primed seedlings exhibited enhanced activation of UPR genes compared to control, whereas unprimed seedlings fail to induce a comparable response following heat stress. (B) Panel depicts representative images of UPR mutants, *bzip60* and *ire1a/b* lines, post-priming/triggering heat stress. Primed *bzip60* and *ire1a/b* knockout mutants survived severe heat stress, whereas the unprimed seedlings failed to survive (n=3 biological replicates). (C-D) Primary root length and fresh weight were quantified at 7 DPS across all genotypes. For WT, primed seedlings showed comparable root length to control, whereas, primed *bzip60* and *ire1a/b* mutants showed a significant reduction in root length compared to their respective controls. Fresh weight was found to be reduced in primed seedlings of all genotypes when compared to their respective control. On the x-axis, C denotes control and P denotes primed seedlings. Data are presented as median ± SD (n=3 biological replicates, comprising a total of 15-24 seedlings represented as individual grey circles). Statistical significance was assessed using one-way ANOVA followed by Tukey’s post-hoc test. Compact letter display was used to denote statistically significant differences among groups (p < 0.05); groups not sharing a common letter are significantly different. (E) To assess the HSR status in *bzip60* and *ire1a/b* mutants, western blot analysis of HSP17.6, HSP21, HSP70, HSP90, and HSP101 proteins was performed for all the conditions post-heat stress. The data revealed that priming led to an increase in the abundance of HSPs in *bzip60* and *ire1a/b*, which persisted even after the stress removal, indicating an active HSR in the mutants. Unprimed *bzip60* and *ire1a/b* seedlings did not show any increase in HSP levels, similar to unprimed wild-type (n=3 biological replicates). [AP - After priming; AT - After triggering (Day 0); DPS - Days post-stress].

To further evaluate whether the loss of UPR impacts priming-mediated responses, we subjected *bzip60* and *ire1a ire1b* (referred to as *ire1a/b*) homozygous mutants to our experimental regime. The IRE1-bZIP60 signalling axis is one of the major branches of plant UPR. Upon ER stress, IRE1 is activated, leading to the formation of *bZIP60s*, which are known to transcriptionally modulate a subset of genes involved in the ERQC/ERAD pathways of the UPR (Nawkar et al., 2018; Pastor-Cantizano et al., 2020). The dynamics of *bZIP60s* accumulation during the experimental regime are shown in **Figure 2B**. When directly exposed to triggering heat stress (unprimed), both *bzip60* and *ire1a/b* seedlings failed to survive, similar to WT (Col-0) (**Figure 3B**). The primed UPR mutants were able to survive severe heat stress upon priming (**Figure 3B**); however, their growth recovery post-stress was found to be slower compared to WT. When observed at seven days post-stress, the root length was found to be significantly lower in the primed *bzip60* and *ire1a/b* compared to their respective controls, while the root length of primed WT was found to be similar to its control (**Figure 3C**). When comparing fresh weight, primed seedlings of all genotypes exhibited lower fresh weights than their respective controls (**Figure 3D**).

As the UPR mutants can survive severe heat stress following priming treatment, we further evaluated whether the loss of UPR influences the HSR. For this, we assessed the chaperone abundance in the *bzip60* and *ire1a/b* mutants. Primed *bzip60* and *ire1a/b* seedlings displayed elevated levels of all the HSPs tested (**Figure 3E**). However, slight differences still existed in the HSP abundance dynamics across genotypes. For instance, HSP17.6 and HSP21 showed similar induction patterns across WT and *ire1a/b*, wherein both the HSPs reached maximal induction by 1 DPS, whereas the *bzip60* mutant displayed a maximum level of HSP21 at 2 DPS (**Figure S4A**). Nonetheless, primed WT showed a slightly higher abundance of both HSP17.6 and HSP21 at 0 DPS compared to *bzip60* and *ire1a/b*. Similarly, both mutants displayed differences in the induction of HSP70 and HSP101 compared to WT. The induction of HSP101 was highest in WT compared to the mutants at 0 DPS post-heat stress, which declined from 0 to 3 DPS. Both the mutants displayed the highest abundance only at 1 DPS, whereas *bzip60* showed a steady decline from 1 to 3 DPS (**Figure S4A**). For HSP70, the level of induction in the mutants and WT was minimal, compared to other HSPs, at all the time points tested (**Figure S4A**). Unprimed *bzip60* and *ire1a/b* displayed no detectable levels of HSPs (**Figure 3E**), similar to the unprimed WT. To assess if the protein level changes parallel the transcript level changes, we determined the expression of *HSPs* in both mutants. As expected, all the *HSPs* tested displayed elevated transcript levels in both primed *bzip60* and *ire1a/b* (**Figure S4B**). Unprimed *bzip60* and *ire1a/b* also displayed a delayed increase in *HSP* transcript levels (**Figure S4B**), similar to WT (**Figure S2D**). Together, these results suggest that, unlike HSR, the UPR signalling is not essential for seeding survival upon priming; however, it may be required for efficient growth recovery post-heat stress.

### Primed seedlings exhibit temporally regulated autophagy

After assessing the dynamics of protein maintenance pathways, we next examined the status of the protein clearance mechanism in response to priming/triggering heat stress. Autophagy, as a clearance pathway, is known to be regulated at both transcriptional and post-translational levels. Transcriptionally, autophagy activation involves the upregulation of autophagy-related genes (ATGs) (Wang et al., 2020; Zhou et al., 2014), whereas at the protein level, autophagy induction is marked by the lipidation of ATG8 to ATG8-PE, a critical step for autophagosome formation (Huang et al., 2022). Upon exposure to priming-stimulus, several *ATGs* were upregulated in the primed seedlings, which was not observed post-triggering stress in both primed and unprimed conditions (**Figure 4A**).

**Figure 4:**
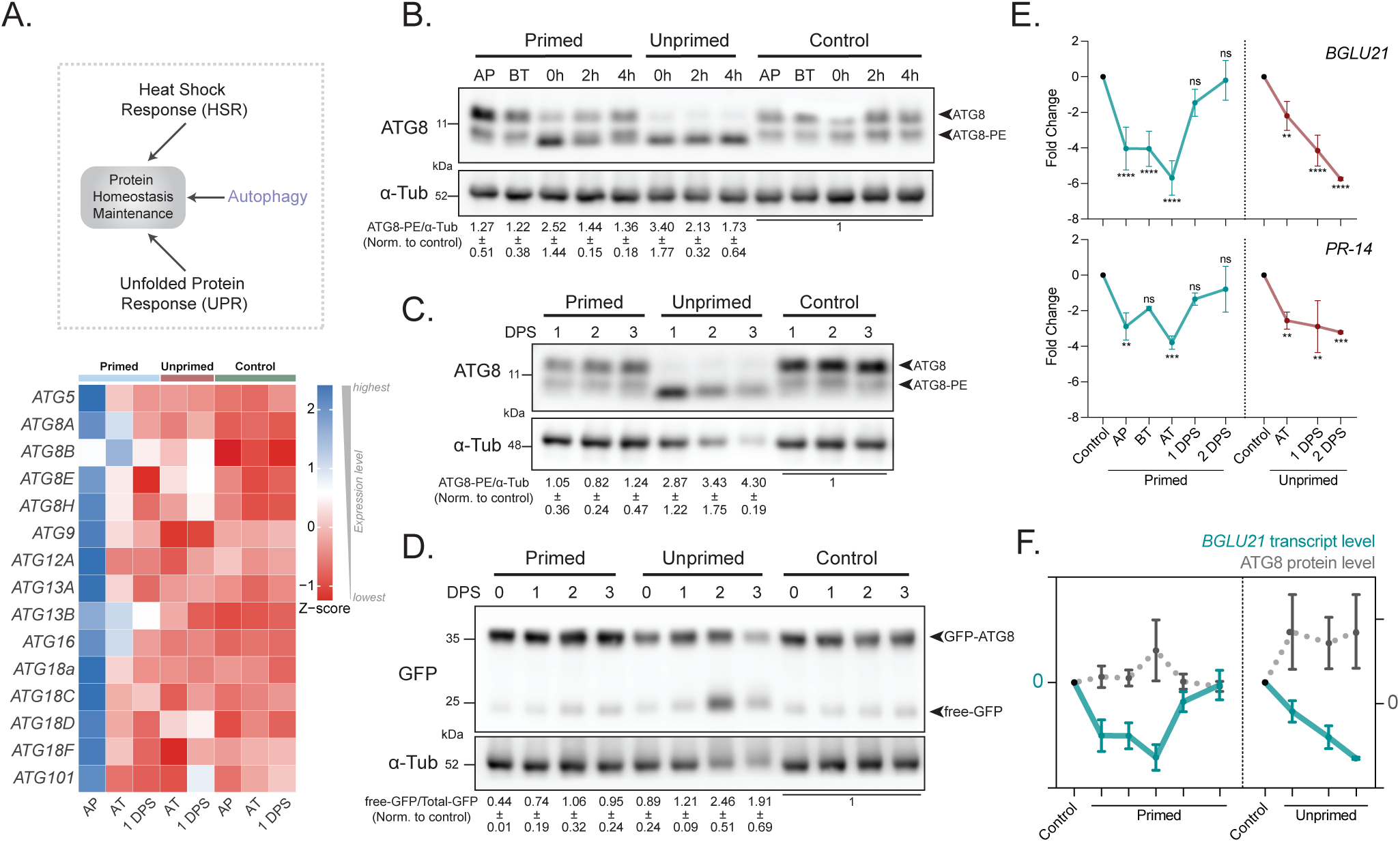
Primed seedlings display temporary autophagy, whereas unprimed seedlings experience sustained autophagy. (A) The heat map shows the mean expression levels of differentially expressed autophagy-related genes (ATGs) across time points, derived from transcriptome analysis. The data indicate that the priming stimulus leads to an increase in the expression of ATGs compared to the control. (B-C) Western blot analysis of ATG8 revealed that primed seedlings showed slightly induced autophagy (as seen by ATG8-PE levels) compared to control at the AP time point, which increased further post-triggering stress (0h to 4h). However, at 1 DPS, autophagy in primed is compara-ble to control levels. In contrast, unprimed seedlings displayed a surge in the ATG8-PE levels post-stress (0h) that remained continuously higher compared to the control. Numbers below denote intensity values post-normalisation with respect to the corresponding control time point (n=3 biological replicates). (D) Western blot of GFP-ATG8 and free-GFP (both detected using anti-GFP antibody) for various samples at the post-heat stress time points revealed that autophagy flux in unprimed seedlings remained high during the stress recovery period, as seen by the high free-GFP levels, whereas autophagy in primed seedlings was similar to that of the control. Numbers below represent normalised mean intensity of free-GFP/Total-GFP (GFP-ATG8+free-GFP) with respect to the corresponding control time point (n=2 biological replicates). (E) Expression of autophagy inhibitors *BGLU21* and *PR-14* is represented as fold change compared to control. In primed, expression of autophagy inhibitors was found to be downregulated at AP, BT and AT timepoints, but it returned to control levels by 1 DPS. However, in unprimed, autophagy inhibitors were continuously maintained at lower levels post-stress. The data are presented as mean ± SD (n=2/3 biological repli-cates). Statistical significance was calculated using one-way ANOVA (**** p <0.0001; *** p <0.001; ** p <0.01; ns - not significant). (F) Data-driven illustrated representation depicting the correlation between the downregulation of the autophagy inhibitor *BGLU21* coinciding with the induction of ATG8 levels in primed and unprimed seedlings. [AP - After priming; BT - Before triggering; AT - After triggering (Day 0); DPS - Days post-stress].

We next quantified the protein level autophagy response by using ATG8 as a marker. We observed that exposure to the priming temperature alone could induce ATG8 levels (**Figure 4B**), as also reflected from transcriptome data. However, immediately after exposure to triggering heat stress (0h), autophagy (ATG8-PE) levels peaked in both primed and unprimed seedlings compared to the control (**Figure 4B**). To assess if the heat stress-induced autophagy sustains, declines or surges with time, we checked the ATG8-PE levels from 1 to 3 DPS. In the unprimed seedlings, the ATG8-PE levels increased further by 1 DPS, which continued until 2 DPS (**Figure 4C**). Notably, the visible decline in ATG8 levels (at 3 DPS) correlated with the reduced protein content in the unprimed seedlings (**Figure S1A**). However, in the primed seedlings, ATG8-PE levels declined by 1 DPS to levels comparable to the control (**Figure 4C**). We also measured the autophagy flux across our paradigm using the GFP-ATG8 processing assay (Kang et al., 2018; Shin et al., 2014). Autophagy flux, as measured by the free-GFP to total-GFP ratio, was high in unprimed seedlings, indicating highly active autophagy compared to primed and control (**Figure 4D**). Such an increase *vs.* decrease in autophagy levels in the unprimed *vs.* primed, respectively, suggests a dynamic regulation of autophagy across these conditions.

### Regulation of autophagy under heat stress is TOR and SnRK-independent

Previous studies have identified the role of TOR and SnRK in the regulation of autophagy under stress (Pu et al., 2017; Soto-Burgos & Bassham, 2017). We wondered if TOR/SnRK are involved in the priming-mediated regulation of autophagy under heat stress. To test this, we analysed the phospho-S6K (pS6K) levels as a measure of TOR kinase activity (Upadhyaya et al., 2020) and active-SnRK (pAMPK: using phospho-AMPKα antibody) for SnRK activity (Baena-González et al., 2007; Belda-Palazón et al., 2020). Interestingly, neither pS6K **(Figure S5, A-B**) nor the pAMPK levels **(Figure S5, C-D**) were altered across all the conditions. These findings suggest that heat stress induced autophagy (as observed in **Figure 4, B-C**) is independent of TOR and SnRK mediated regulation.

### Priming-mediated autophagy induction is associated with attenuation of autophagy inhibitors

The UPR regulator IRE1 has also been linked to modulate autophagy by fine-tuning its RIDD (Regulated IRE1-dependent decay of mRNA) activity (Li & Howell, 2021; Maurel et al., 2014). Notably, RIDD leads to the downregulation of inhibitors of autophagy, such as *BGLU21*, *ROSY1* and *PR-14,* by targeting those mRNAs for degradation (Bao et al., 2018). We surmised that reversal to basal level *vs.* more sustained autophagy in the primed *vs.* unprimed, respectively, may involve RIDD-mediated regulation. Therefore, we examined the steady state transcript levels of *BGLU21* and *PR-14* across all the conditions. At the time points-post priming (AP), before triggering (BT) and after triggering (AT: 0 DPS), primed seedlings showed a significant reduction in the transcript levels of both the genes compared to the control (**Figure 4E**). However, by 1-2 DPS, we observed a recovery in their transcript abundance (**Figure 4E**). In contrast, the transcript levels of *BGLU21* and *PR14* remained significantly lower in unprimed seedlings at all the time points compared to the control (**Figure 4E**). In particular, these changes in the transcript abundance correlated with the ATG8/ATG8-PE levels in these seedlings. For instance, in the primed, reduced transcript abundance of these autophagy inhibitors correlates with the induced autophagy (at AP, BT and AT). By 1-2 DPS, when the transcript abundance of these inhibitors reverted to near control level, autophagy was found to return to basal level in the primed (**Figure 4F**). Contrarily, sustained induction of autophagy aligns with reduced transcript abundance of the autophagy inhibitors in the unprimed, post-exposure to triggering heat stress (**Figure 4F**). These observations suggest that priming modulates autophagy, potentially through the RIDD-mediated attenuation of autophagy inhibitors.

Further, we assessed whether the autophagy dynamics are affected in the IRE1A/B, and its downstream target, bZIP60, knockout mutants. Interestingly, we observed that autophagy remained induced in both *bzip60* and *ire1a/b* mutants (**Figure S6, A-B**) and its dynamics were similar to those of WT (**Figure 4, B-C**). Loss of IRE1A/B and bZIP60 did not seem to disrupt autophagy induction during heat stress. This was surprising, as we observed that heat stress facilitated IRE1-dependent dynamic degradation of autophagy inhibitors, thereby modulating autophagy. However, autophagy was still being induced in *bzip60* and *ire1a/b* mutants. These findings suggest the involvement of additional IRE1-independent mechanism that may operate during heat stress to regulate autophagy.

### Impairment of the protein protective response amplifies the protein clearance mechanisms

Given the potential involvement of alternate mechanisms regulating autophagy, we next investigated the role of HSFA1a, a transcription factor previously shown to induce autophagy under abiotic stress conditions (Wang et al., 2015; Xie et al., 2022). Notably, we observed the induction of HSFA1 activity and a parallel upregulation of *ATG* genes post-priming stimulus (**Figure 4A; S2E**). When we probed to assess if autophagy response gets affected in the *hsfa1a/b/d* triple mutant, we observed that autophagy remains active in both primed and unprimed as evidenced by the sustained accumulation of ATG8-PE post-stress (**Figure 5A**), suggesting that the autophagy response may be under a broader multiple regulatory system.

**Figure 5:**
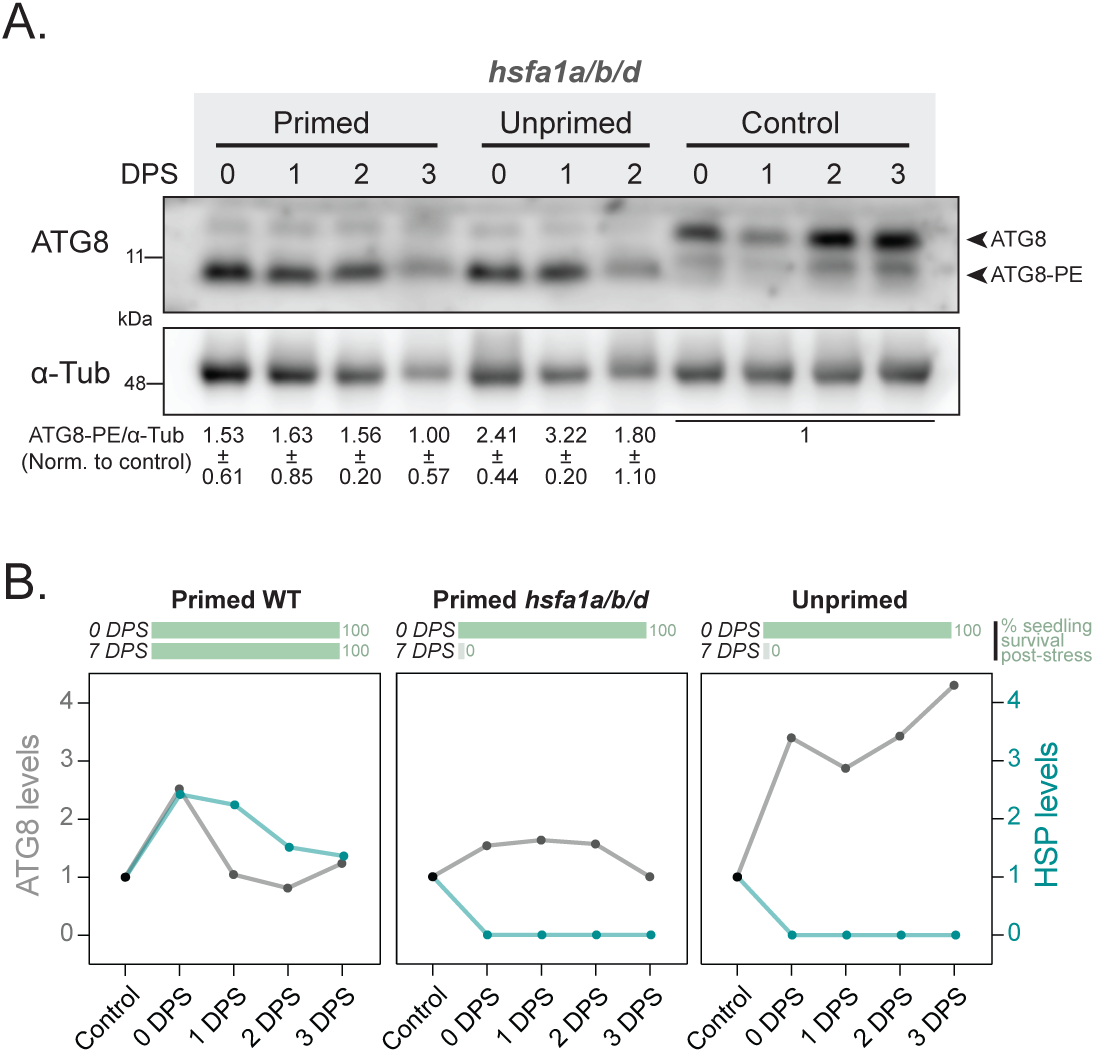
Impairment of the heat shock response amplifies autophagy during proteotoxic stress. (A) Western blot analysis of ATG8 in *hsfa1a/b/d* mutants shows that autophagy remains highly induced in both primed and unprimed following heat stress (n=3 biological replicates). The numbers below denote intensity values post-normalisation with Data-driven illustrated line plots comparing ATG8 and HSP abundance profiles in primed WT, primed *hsfa1a/b/d*, and unprimed seedlings at the indicated timepoints. The bar plots above indicate the percent-age of seedling survival at 7 DPS (n=3 biological replicates per condition). In primed WT, chaperone accumulation (HSPs) peaks early and is accompa-nied by transient ATG8 induction, suggesting efficient proteostasis and resolution of stress via timely atten-uation of autophagy. Conversely, both unprimed WT and primed *hsfa1a/b/d* mutants failed to induce HSPs, correlating with sustained ATG8 accumulation and complete post-stress mortality. Collectively, these data illustrate that thermotolerance is driven by a coordinated HSP-autophagy interplay, whereas its failure leads to autophagy overactivation and seed-ling lethality. [DPS - Days post stress].

As we observed that the HSP induction was abolished in the *hsfa1a/b/d* mutant with or without priming (**Figure 2J**), we accounted for all the protein level evidence and summarized it in a data-directed illustration (**Figure 5B**). Collectively, as evidenced from primed WT, activation of protein protective response (chaperone accumulation) facilitates a dynamic clearance response (autophagy) that subsides during stress recovery (**Figure 5B**). However, as observed in unprimed (all genotypes) and primed *hsfa1a/b/d* seedlings, failure to accumulate chaperones upon heat stress may facilitate the persistent activation of autophagy, aimed to resolve stress, but ultimately leads to lethality (**Figure 5B**).

### Unprimed seedlings accumulate protein aggregates following triggering heat stress

So far, our findings indicate that priming-mediated heat stress tolerance involves the accumulation of chaperones and temporary activation of autophagy. To test whether these changes contribute to the effective maintenance of protein homeostasis, we examined the proteostasis state by analysing protein aggregates at 1 DPS, the time at which we observed differential autophagy between the primed and unprimed conditions (**Figure 4C**). The unprimed seedlings displayed elevated ubiquitin-conjugated aggregates compared to primed or control (**Figure 6A**). Additionally, size distribution analysis of the aggregates using dynamic light scattering (DLS) clearly showed that the unprimed seedlings accumulated a high proportion of larger aggregates (13.03nm, 460.1nm and 6662.6nm), whereas the primed seedlings accumulated comparatively smaller aggregates (18.02nm, 174.1nm and 1977.0nm), which were proportionally similar to control (24.9nm, 136.5nm and 1550.5nm) (**Figure 6B**). We also compared the diffusion behavior of the aggregates by using the DLS data. BSA was used as a standard monomeric protein control (8nm), and it exhibited the fastest (steepest) decay (**Figure 6C**). The aggregates derived from the unprimed displayed the slowest decay, reflecting larger population sizes, whereas the aggregates derived from the primed showed a comparatively faster decay, reflecting a relatively smaller population (**Figure 6C**). The control samples showed a profile comparable to BSA. Taken together, our study indicates that thermopriming facilitates timely activation of the heat shock response, unfolded protein response and autophagy, thereby aiding in effective aggregate clearance, and collectively promotes heat stress tolerance.

**Figure 6:**
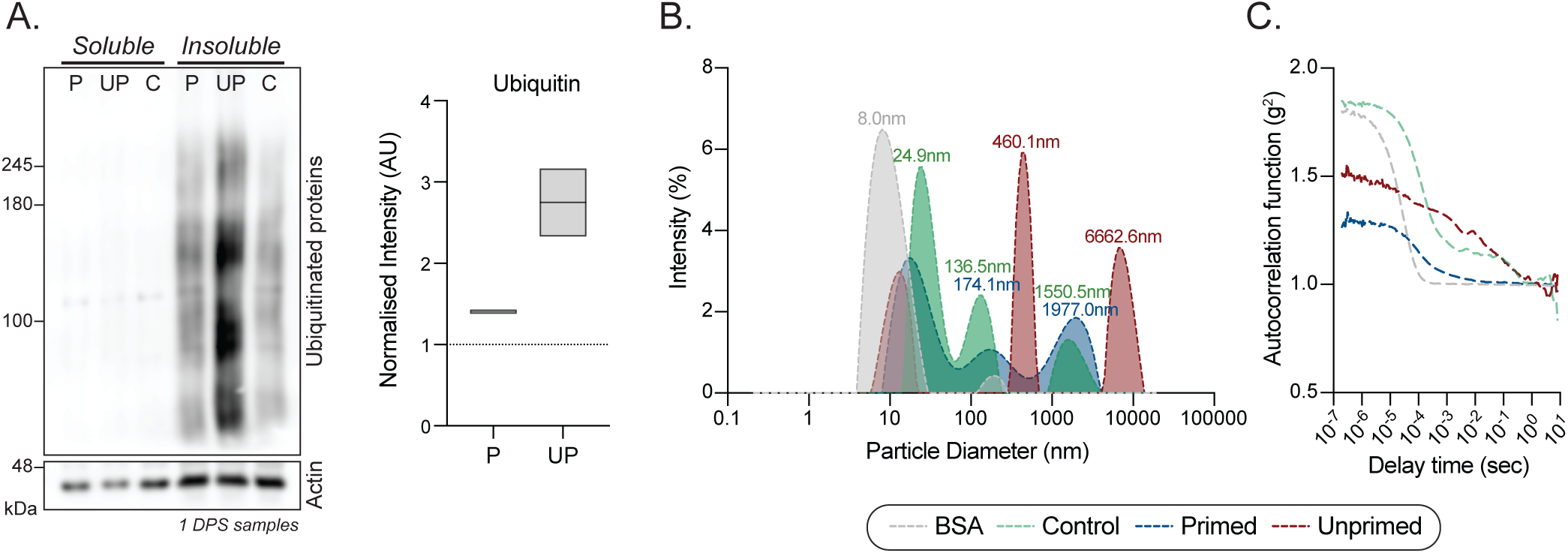
Priming facilitates effective aggregate clearance following heat stress. (A) Immunoblot analysis of ubiquitin-conjugated aggregates shows that unprimed seedlings have a higher accumulation of aggregates compared to primed and control seedlings by 1 DPS. Box plot represents the normalised mean intensity for ubiquitin in the insoluble fraction. The dotted line denotes the control value (n=2 biological replicates). (B) Dynamic Light Scattering (DLS) analysis showing the particle size distribution of protein aggregates isolated at 1 DPS from control, primed, and unprimed seedlings. Bovine serum albumin (BSA) was used as a monomeric protein standard. Unprimed seedlings accumulated a higher proportion of large aggregates (460.1 nm and 6662.6 nm), indicating extensive aggregation. In contrast, primed seedlings displayed smaller aggregate populations (174.1 nm and 1977.0 nm), comparable to control, suggesting more efficient aggregate clearance. (C) Autocorrelation function (g²) curves derived from DLS measurements reflect particle size dynamics and diffusion behavior. BSA exhibited the steepest/fastest decay, consistent with uniform small particles. Aggregates derived from unprimed seedlings exhibited the slowest decay, reflective of larger, less diffusive aggregate populations. Primed samples show a faster decay than unprimed, indicating smaller and more dynamic aggregates. Data is representative of two biological replicates.

## Discussion

In the current study, we demonstrated that the efficient regulation of protein homeostasis underlies priming-mediated short-term acquired thermotolerance in Arabidopsis. Heat stress has a profound impact on the thermolabile molecular complexes, resulting in protein dysfunction and aggregation (McLoughlin et al., 2019; Mittler et al., 2012; Rosenzweig et al., 2019). Maintaining proteostasis under stress requires a fine balance between timely refolding of proteins via robust chaperone activity, and the efficient clearance of irreversibly damaged proteins through the action of disaggregases, autophagy, and/or the UPS (Hartl et al., 2011; McLoughlin et al., 2019; Mogk et al., 2018). Disruption of this balance can lead to proteotoxic stress, which ultimately compromises cellular viability. In this study, we observed that direct exposure to heat stress severely affected the protein content in the unprimed seedlings, which was alleviated in the primed seedlings. These observations are consistent with the previous reports showing that protein content is reduced upon exposure to noxious heat stress (Gulen & Eris, 2004; Nagy-Réder et al., 2022). We found that thermopriming facilitates an increase in chaperone abundance and UPR activation along with a temporary autophagy response, which together contribute to the maintenance of protein homeostasis.

The unprimed seedlings, even upon facing similar triggering heat stress, failed to accumulate HSPs despite a significant expression at the transcript level. Such a failure to accumulate HSPs may stem from HSP sequestration within aggregates or impaired translation post-stress. We observed that genes involved in translation remained severely downregulated in unprimed seedlings compared to primed post-heat stress. Additionally, HSP101, a critical disaggregase that coordinates with other HSPs to resolve proteotoxic stress (McLoughlin et al., 2019), was also detected only in primed seedlings (**Figure S3B**). Cumulatively, this suggests that the chaperone synthesis was completely impaired in the unprimed seedlings. In addition, the unprimed seedlings displayed high levels of ubiquitinylated insoluble proteins, a trend also reported in HSP101 deficient plants during heat stress (McLoughlin et al., 2019). Similarly, Dannfald et al. (2025) showed that priming protects the global translational potential under heat stress, whereas direct heat exposure leads to preferential decay of mRNAs.

Given the difference in autophagy response between primed and unprimed, we examined the role of TOR/SnRK-dependent pathways, which are known to be involved in regulating autophagy (Signorelli et al., 2019; Soto-Burgos & Bassham, 2017; Yang et al., 2023), However, we did not detect any changes in TOR/SnRK activity under our experimental regime. Indeed, TOR-independent mechanisms have been reported to modulate autophagy under oxidative and ER stress conditions (Pu et al., 2017). Since we observed that thermopriming induced ER stress and subsequent UPR activation, we explored the potential involvement of IRE1. IRE1-RIDD activity has been previously associated with autophagy induction under tunicamycin treatment (ER-stress inducing agent), wherein RIDD-mediated degradation of autophagy inhibitors was shown to promote autophagy (Bao et al., 2018). The findings from the current study show that thermopriming facilitates a degradation of RIDD target transcripts, which correlates with the induction of autophagy in primed seedlings. Similarly, in unprimed seedlings, the prolonged autophagy induction was associated with sustained transcriptional downregulation of autophagy inhibitors. While these observations suggest that RIDD activity influences autophagy by modulating the abundance of autophagy inhibitors, the precise mechanism remains to be explored. Bao et al. (2018) suggested that BGLU21 (β-glucosidase 21) and PR-14 (pathogenesis-related protein 14) proteins might interfere with the lipidation of ATG8, a crucial step for autophagosome formation, thereby affecting autophagy.

Notably, despite the loss of IRE1A/B or bZIP60, autophagy was still induced post-heat stress. Our observations concur with the previous report that shows bZIP60 is dispensable for IRE1-dependent autophagy induction during ER stress (Bao et al., 2018). Moreover, although autophagy was found to be induced in *ire1b* mutants following 42°C heat treatment (Yang et al., 2016), autophagy was not induced in *ire1b* and *ire1a/b* mutants under ER stress upon DTT or tunicamycin treatment (Bao et al., 2018; Liu et al., 2012). Given that autophagy was still activated in *ire1a/b* following thermal priming, we posit that alternative pathways may mediate heat-induced autophagy. When probed for the involvement of a heat inducible transcriptional activator of autophagy, *viz.* HSFA1 (Löchli et al., 2023; Wang et al., 2015; Xie et al., 2022), autophagy was found to be induced even in the *hsfa1a/b/d* mutant. These observations suggest that the autophagy response involves a multilayered regulation, a feature advantageous for sessile organisms to tackle challenging environments.

Previous studies have also implicated the involvement of NBR1 (next-to-BRCA1) and ATG8-interacting protein 1 (ATI1), 2 (ATI2) and 3 (ATI3) in mediating selective autophagy (Jung et al., 2020; Sjøgaard et al., 2019; Thirumalaikumar et al., 2020; Zhou et al., 2018). The autophagy response can either be selective, which involves recognition by specific receptors followed by subsequent clearance of damaged organelles/aggregates; or bulk, which facilitates the clearance of cytoplasmic debris in a non-selective manner (Marshall & Vierstra, 2018). From our transcriptome data, we observed that selective autophagy receptors such as NBR1 and ATI1/2/3A were upregulated in the primed seedlings alone (**Figure S7**), suggesting that the observed autophagy response in these seedlings may potentially be selective in nature. Contrastingly, bulk autophagy may operate in the unprimed, as we did not observe any change in the expression of these receptors. Moreover, vacuolar cell death has been previously correlated with high autophagic flux (Minina et al., 2013), a feature we observed in unprimed seedlings. Given the present evidence, it is tempting to speculate that bulk autophagy drives the unprimed seedlings towards vacuolar cell death, however, this requires further validation.

Additionally, analysis of the *bzip60* and *ire1a/b* knockout mutants under primed conditions revealed reduced growth recovery post-stress compared to WT. Previous studies have reported that *bzip60* and *ire1a/b* mutants exhibit severe growth retardation under chemically-induced ER stress conditions (Deng et al., 2013; Pu et al., 2019). Primed *bzip60* and *ire1a/b* seedlings displayed an increase in HSP levels; however, their response was delayed compared to WT. One possible explanation is attenuated/delayed protein translation, a feature previously reported in *ire1a/b* mutants under ER stress (Yoo et al., 2024). Notably, Yoo et al. (2024) also demonstrated that supplementation with synthetic chaperones mitigated the effects of attenuated translation and growth retardation in *ire1a/b* mutants. Given that the *ire1a/b* mutants accumulate chaperones during the tested regime, the priming stimulus facilitates a faster and more robust recovery of these seedlings under stress. Taking together the observed altered dynamics of HSP induction in UPR mutants, we suggest a potential crosstalk between the HSR and UPR pathways during thermotolerance. We also observed that the *hsfa1a/b/d* mutant failed to survive heat stress even upon exposure to a priming stimulus. Such a failure could arise due to the inability of the *hsfa1a/b/d* mutant to induce HSPs, implying that efficient protein maintenance (by chaperones) is a critical determinant of stress response outcomes. An earlier report suggested that autophagy contributes to the degradation of HSP, as autophagy mutants showed prolonged chaperone accumulation (Sedaghatmehr et al., 2019). However, our findings suggests that autophagy and chaperone response may act cooperatively to modulate protein homeostasis, with the impairment of one pathway potentially amplifying the activity of the other, as seen in *hsfa1a/b/d* where failure to induce HSPs is accompanied by sustained autophagy. These evidences underscores the importance of maintaining a delicate balance between protein -folding and -clearance machineries in restoring cellular homeostasis following stress. The unprimed seedlings experience a collapse of this balance, as evidenced by the failure to accumulate HSPs, lack of UPR activation, excessive aggregate buildup, and sustained autophagy, which ultimately compromises their growth and survival (**Figure 7**).

**Figure 7:**
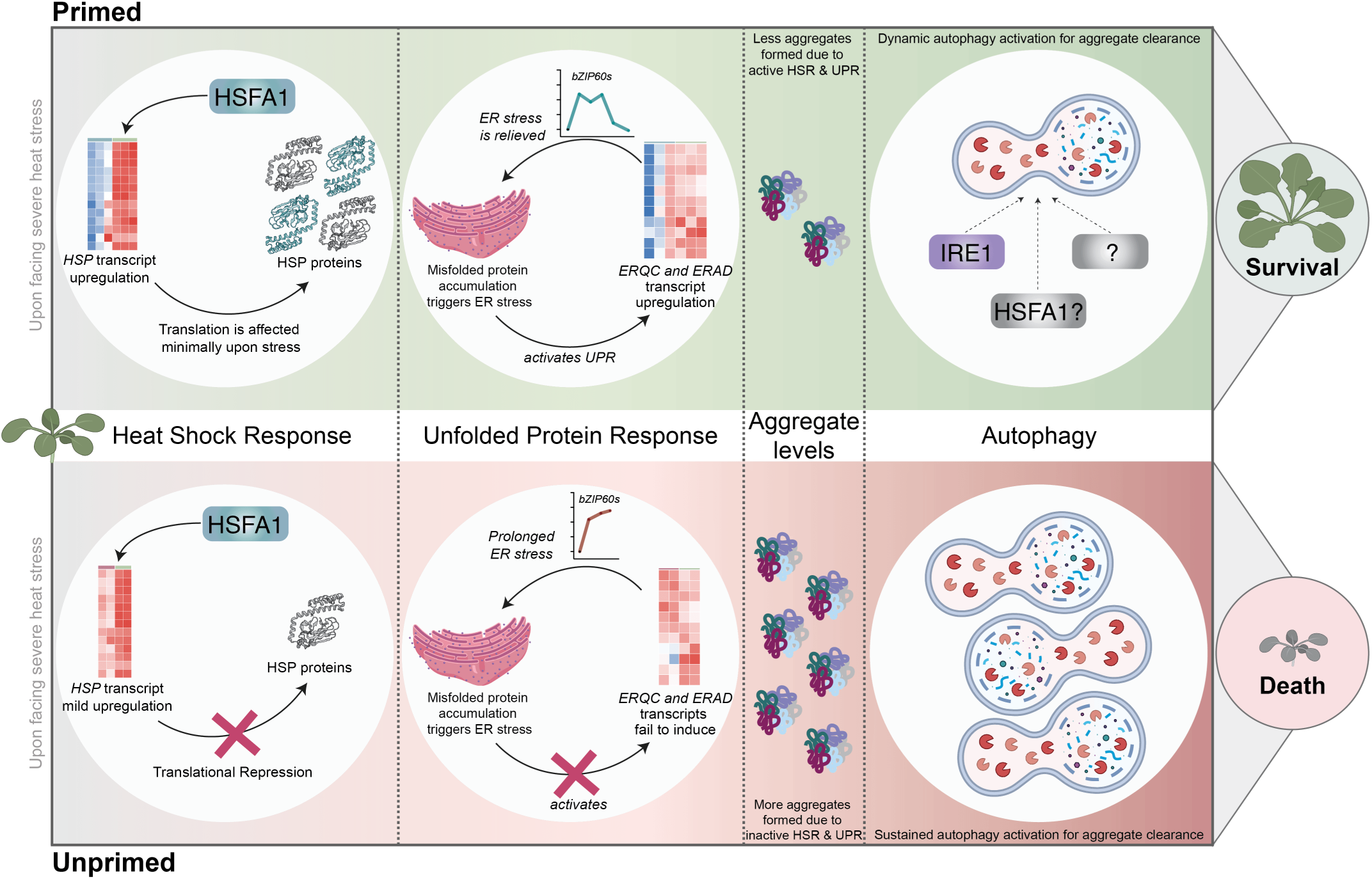
Priming-mediated acquired thermotolerance underlies dynamic regulation of proteostasis. The illustration compares the responses of primed (top panel) and unprimed (bottom panel) Arabidopsis seedlings upon exposure to severe heat stress. In primed seedlings, HSFA1-mediated transcriptional activation drives a robust heat shock response (HSR) associated with accumulation of HSPs. Accumulation of misfolded proteins in the endoplasmic reticulum (ER) activates the unfolded protein response (UPR), leading to induction of transcripts involved in ER-quality control (ERQC) and ER-associated degradation (ERAD) pathways, thereby alleviating ER stress. As a result of active HSR and UPR, aggregate formation is restricted (minimal). Short-term autophagy, potentially regulated by IRE1 -dependent and -independent pathways, efficiently clears residual aggregates, maintaining proteostasis and promoting survival. In contrast, unprimed seedlings undergo translational repression and fail to efficiently activate HSR and UPR, resulting in prolonged ER stress and excessive protein aggregation. In response, autophagy remains persistently activated but soon becomes potentially maladaptive, coupled with a proteostasis collapse and ultimately leading to seedling death. *Some elements of the illustration were taken from BioRender.com*.

Taken together, our study highlights that thermopriming facilitates the efficient maintenance of protein homeostasis through the timely activation of HSR, UPR and autophagy, thereby aiding in seedling survival upon experiencing severe heat stress. Furthermore, we identified that UPR activation is important for timely seedling recovery post-stress, but it may not be essential for mounting thermotolerance. However, HSFA1-mediated HSR is indispensable for mounting a heat stress tolerance response. As we observe chaperone accumulation is affected in UPR mutants, further studies are required to understand the probable crosstalk between UPR and HSR pathways during thermotolerance. Additionally, how heat stress may facilitate the mounting of bulk vs. selective autophagy during acquired thermotolerance needs to be further understood. Collectively, our findings revealed that precise and coordinated regulation of protein folding and clearance pathways is a key determinant of thermotolerance in Arabidopsis.

## Methods

### Plant materials and growth conditions

Arabidopsis (Col-0-referred to as WT and Ws-0-referred to as Ws-0) seeds were surface sterilized by washing with 80% ethanol for 30 seconds (s) followed by 1% bleach for 3 minutes (min), and 3x washes with sterile double distilled water. Seeds were then sown on ½ Murashige and Skoog media with 1% sucrose and 0.8% agar powder. The seeds were stratified at 4°C for two days before being transferred to growth chambers with 16-hour (h) light and 8 h dark, 65% humidity. The five-day-old Arabidopsis seedlings were primed by exposing them to 37°C for 90 min, followed by a recovery at 22°C for 90 min, and heat-triggering at 45°C for 45 min. The unprimed seedlings were directly exposed to 45°C for 45 min. Control seedlings were maintained under optimal growth conditions (22°C, 16h light and 8h dark, 65% humidity) throughout the experiment. Seedlings were harvested at different stress recovery time points for gene expression and protein analysis. The treatment conditions described above were also used for the analysis of GFP-ATG8a (CS39996) and *bzip60* (CS813074) (both lines were obtained from ABRC (<u>A</u>rabidopsis <u>B</u>iological <u>R</u>esource <u>C</u>enter)). The *ire1a/b* line was kindly obtained from Chen et al., (2013) and Pu et al., (2019). Homozygous *hsfa1a/b/d* (psi00284) line was obtained from RIKEN-BRC.

### Physiological measurements

To determine the effects of priming or triggering heat stress on growth, the seedlings were analysed for leaf number, root length and fresh weight. Data on leaf number (48 seedlings from three independent experiments), primary root length (18 seedlings from three independent experiments) and fresh weight (three biological replicates) were collected and analysed. For Figure 1E, 16 seedlings were pooled to form a single replicate, and for Figure 3D, individual seedling fresh weight was measured. For leaf number measurements, both true leaves and cotyledons were considered. Root length was measured using the ImageJ software (https://imagej.net/software/fiji/) (Schneider et al., 2012).

### RNA extraction and gene expression analysis

For extracting RNA, approximately 40-50 mg of Arabidopsis seedlings were collected at several time points (post-priming and triggering, and during recovery) and frozen in liquid nitrogen. The samples were ground using Retsch Miller MM400 at 25Hz for 30 s (x2 cycles). After grinding, RNA was isolated from the samples using a combination of TRIzol/chloroform extraction and kit method (PureLink™ RNA Mini Kit, Thermo; 12183018A). The purified RNA was treated with DNase I (NEB; M0303) following standard protocol. cDNA was synthesized using 1 µg of RNA following the manufacturer’s instructions (1st strand cDNA synthesis kit, Takara; 6110A). For gene expression analysis, RT-PCR was performed using SapphireAmp® Fast PCR Master Mix (Takara; RR350A), while iTaq™ Universal SYBR® Green Supermix (Biorad; 1725121) was used for performing quantitative real-time-PCR. The housekeeping gene *ACT2* was used as an internal control to normalize the target gene expression across samples. All the primers used along with their corresponding gene accession ID are described in **Table S1**.

Transcriptome analysis was performed with two biological replicates under the described conditions (**Figure S2A**). Paired-end sequencing libraries were prepared using SMART-Seq Library Prep Kit (R40047) as per the manufacturer’s protocol and the sequencing was done using Illumina Novaseq 6000 S4 platform. Adapter trimming and quality control was performed using FASTP (Chen et al., 2018) and FASTQC (https://www.bioinformatics.babraham.ac.uk/projects/fastqc/) respectively. Kallisto v.0.50.1 (Bray et al., 2016) was used for indexing of the reference Arabidopsis transcriptome downloaded from Ensembl. Read mapping was performed using Kallisto following which TxImport (Soneson et al., 2016) was used to read the Kallisto outputs in the R environment. Differential gene expression analysis between control, primed and unprimed seedlings was performed using the edgeR package (Chen et al., 2025) in RStudio. Genes with an adjusted P-value < 0.01 and a cut-off of Log2FC ≥ 1 were used for selecting upregulated genes, and Log2FC ≤ −1 was used for selecting downregulated genes. The expression profiles of the DEGs used for plotting are available as **Dataset S1**.

### Protein extraction and Western blot analysis

Soluble protein was extracted from Arabidopsis seedlings as previously described by Yamamoto et al. (2018) with a few modifications. About 40-50 mg of seedlings was collected and frozen in liquid nitrogen. The samples were ground using a Retsch Miller MM400 at 25Hz for 30 seconds (s) (x2 cycles). After grinding, 200-300µl of 1x RIPA lysis buffer (Merck; 20-188) containing 10% glycerol, 1x phosphatase inhibitor cocktail (PhosSTOP, Roche; 4906837001), and 1x protease inhibitor cocktail (Roche; 4693132001) was added and incubated at 4°C for 1 h at 1500 rpm. The samples were centrifuged at 14000 rpm for 30 min at 4°C, after which the supernatant containing protein was collected and stored at -80°C until further use. Plant protein aggregates were extracted using a standard protocol as described by Phukan et al. (2023).

For protein analysis, the protein concentration was estimated using the BCA method, following the manufacturer’s instructions (Cat. No. 23225). Protein (20 µg per lane) was loaded onto an 8% SDS-PAGE gel, which was stained with Coomassie Brilliant Blue. Protein bands were visualized after destaining the gel following a standard protocol (“Destaining Solution for Coomassie Brilliant Blue R250,” 2007). For Western blot analysis, 20 µg of protein was loaded onto 6-12% SDS-PAGE, following which the proteins were transferred to the nitrocellulose membrane. The blot was blocked using 5% BSA or 2% skim milk solution for 1 h at room temperature (RT), followed by primary antibody incubation (overnight at 4°C). The blot was developed after incubation with secondary antibody (1 h at RT) using Immobilon Forte HRP substrate (Cat. No. WBLUF0500). α-Tub (CST 3873), ATG8 (Abcam, ab77003), ATG8 (Agrisera, AS14 2769), GFP (CST 2956), HSP70 (Abcam, ab2787), HSP90 (Sigma, CA1023), HSP90 (Agrisera, AS08 346), HSP17.6 (Agrisera, AS07 254), HSP21 (Agrisera, AS08 285), HSP101 (Agrisera, AS08 287), pS6K (CST 9234), S6K (CST 9202), pAMPK-α (CST 2535), Ubiquitin (CST 3933) and Actin (AS13 2640) were used as primary antibodies. Anti-rabbit (sc-2313; CST 7074) and anti-mouse (CST 7076) were used as secondary antibodies. Blots were quantified densitometrically using the ImageJ software.

### Dynamic Light Scattering Measurements

To assess the size distribution profiles, 20 µg of aggregate sample or standard was diluted appropriately in ultrapure water and analyzed using the Litesizer 500 system equipped with a BM10 module (Anton Paar Instruments). Measurements were acquired at 25 °C, using a disposable cuvette and side-scatter detection angle in the automatic mode, with a maximum of 60 runs per sample and 10 s per run.

### Statistical Analysis

Data were plotted using GraphPad Prism 10.0 (https://www.graphpad.com/). The experiments were conducted using three biological replicates unless otherwise stated. The data was analysed using One-way ANOVA (for qPCR and phenotype analysis) and Two-way ANOVA (for phenotype analysis) to determine statistical significance. *p* values <0.05 were considered significant (**Dataset S2 and S3**).

## Data Availability

The transcriptome data generated in this study have been deposited to the Sequence Read Archive (SRA) of the National Centre for Biotechnology Information (NCBI) under BioProject ID PRJNA1312973.

## Acknowledgements

MB acknowledges the Council of Scientific and Industrial Research (CSIR), Government of India, for his fellowship. This work was supported by funding from the Department of Science and Technology (Sir JC Bose Fellowship - for BJR), the Science and Engineering Research Board (SPG/2021/004000), and intramural support from IISER Tirupati (ADA and BJR), University of Hyderabad (BJR). We express our sincere gratitude to Prof. Diane Bassham (Iowa State University) and Prof. Federica Brandizzi (Michigan State University) for providing us with seeds of *ire1a ire1b* mutant.

## Authors Contribution

**MB:** Designing the experiments, performing the experiments, interpretation of results, preparation of the figures, writing the manuscript; **ADA:** conceptualization of the research question, supervision of the work, design of the experiments, interpretation of results, writing and editing the manuscript, funding acquisition; **BJR:** conceptualization of the research question, supervision of the work, design of the experiments, interpretation of results, writing and editing the manuscript, funding acquisition.

---

**Figure S1:**
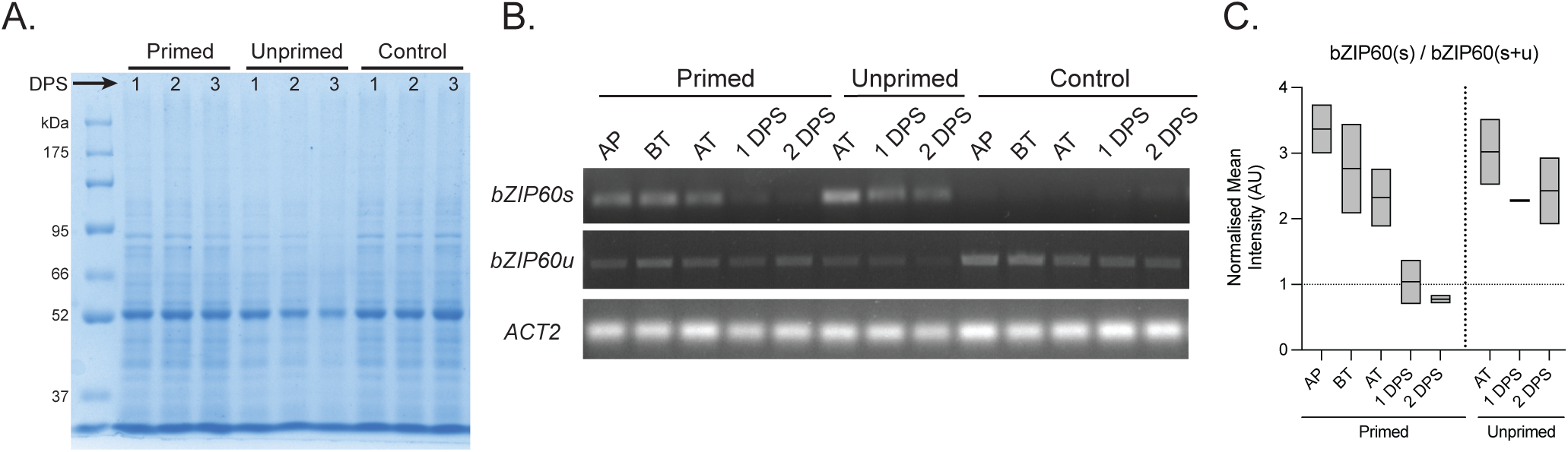
Unprimed seedlings display total protein loss and sustained presence of misfolded proteins following heat stress. (A) SDS-PAGE analysis indicates that unprimed seedlings undergo a reduction in total protein content by the end of 3 DPS, compared to primed and control (n=3 biological replicates). (B-C) The *bZIP60* spliced form (*bZIP60s*) was used as a marker to detect misfolded protein accumulation. RT-PCR of *bZIP60s*, *bZIP60u* and *ACT2* for all the conditions at given time points post heat stress suggested that primed seedlings initially accumulate misfolded proteins but recovered by 1 DPS to control levels (denoted by dotted line). However, unprimed seedlings experienced sustained accumulation of misfolded proteins as seen from the presence of high *bZIP60s*. Box plot represents quantification for *bZIP60*(s) / *bZIP60*(s+u(unspliced)) (n=2 biological replicates). [AP - After priming; BT - Before triggering; AT - After triggering (Day 0); DPS - Days post-stress].

**Figure S2:**
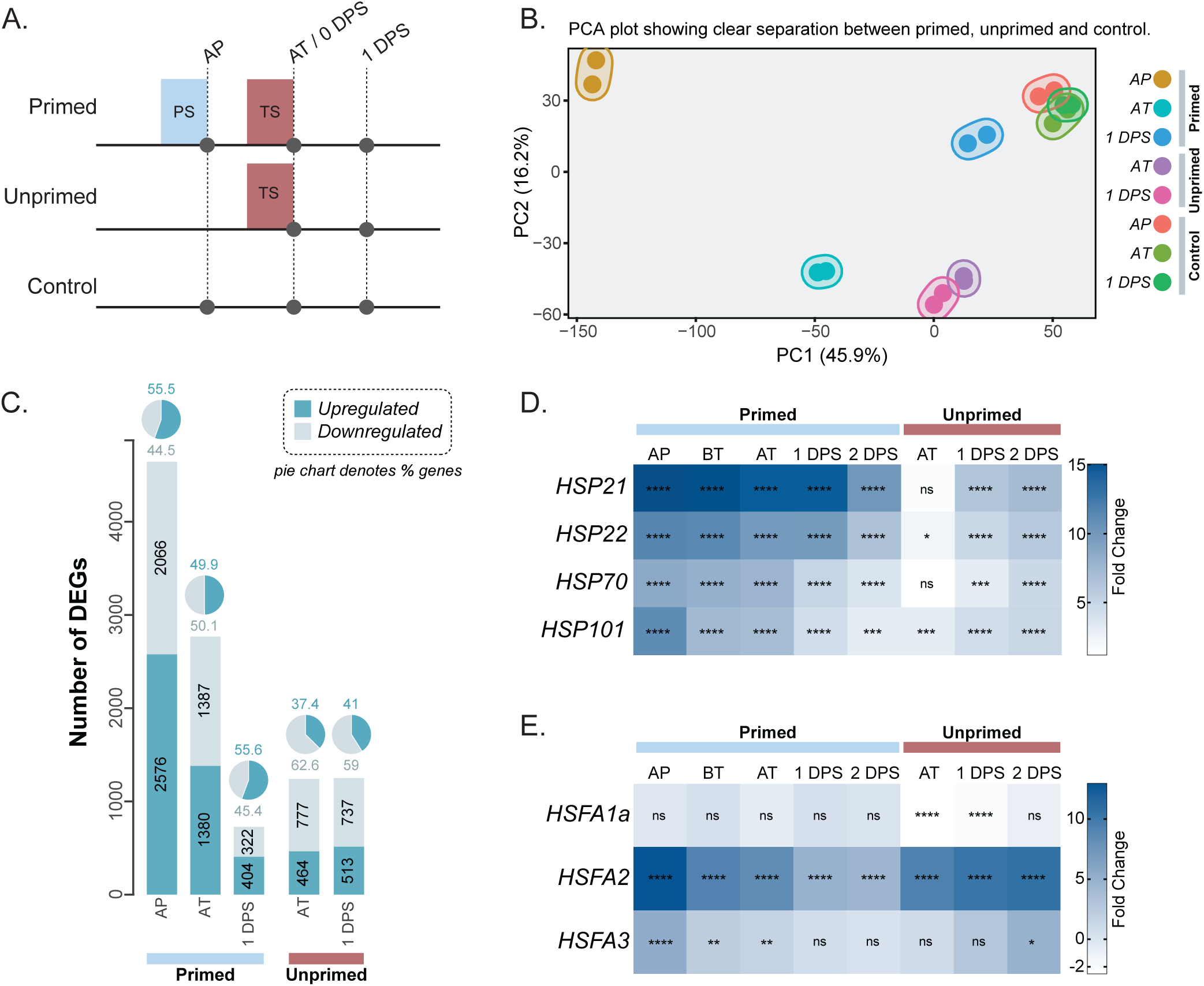
Transcriptome and expression analysis of primed and unprimed seedlings post heat stress. (A) Schematic representation of sampling time points for transcriptome analysis. Seedlings was exposed to a priming stimulus (PS) followed by a triggering stimulus (TS) (primed), only a TS (unprimed), or no treatment (control). Samples were collected after priming (AP), immediately after triggering (AT; 0 DPS), and one day post-stress (1 DPS) (n=2 biological replicates). (B) Principal component analysis (PCA) of transcriptome profiles shows distinct transcriptional reprogramming induced by priming, with AP_primed samples separating strongly from unprimed and control. Upon triggering and during recovery, AT_primed and 1 DPS_primed shift closer to unprimed and control respectively. Unprimed (AT and 1 DPS) and control (AP, AT and 1 DPS) samples form a tighter cluster, indicating transcriptional similarity across samples. (C) Bar plot shows differentially expressed genes (DEGs) in primed and unprimed seedlings (n=2 biological replicates). A total of 6727 unique DEGs were identified with thresholds of log2 fold change ≥ 1 or ≤ –1 and a false discovery rate (FDR) < 0.01. The pie charts on top indicate the percentage of upregulated or down-regulated DEGs for each condition. (D) The heatmap represents mean gene expression levels post-priming/triggering heat stress. Expression of *HSP21, HSP22, HSP70* and *HSP101* reflects fold change compared to control. In the primed, *HSPs* get highly induced upon priming and triggering compared to the control. In contrast, in the unprimed, induced expression for most of the *HSPs* is observed in only 1 DPS. The data are presented as mean ± SD (n=3 biological replicates). (E) Heatmap represents mean gene expression levels of *HSFA1a, HSFA2* and *HSFA3* post-priming/triggering heat stress. *HSFA1a* expression (except AT/1 DPS in unprimed) did not significantly change across time points, whereas *HSFA2* and *HSFA3* expression significantly increased immediately after priming (n=3 biological replicates). Statistical significance was calculated using one-way ANOVA (**** p <0.0001; *** p <0.001; ** p <0.01; * p <0.05; ns - not significant). [DPS - Days post-stress].

**Figure S3:**
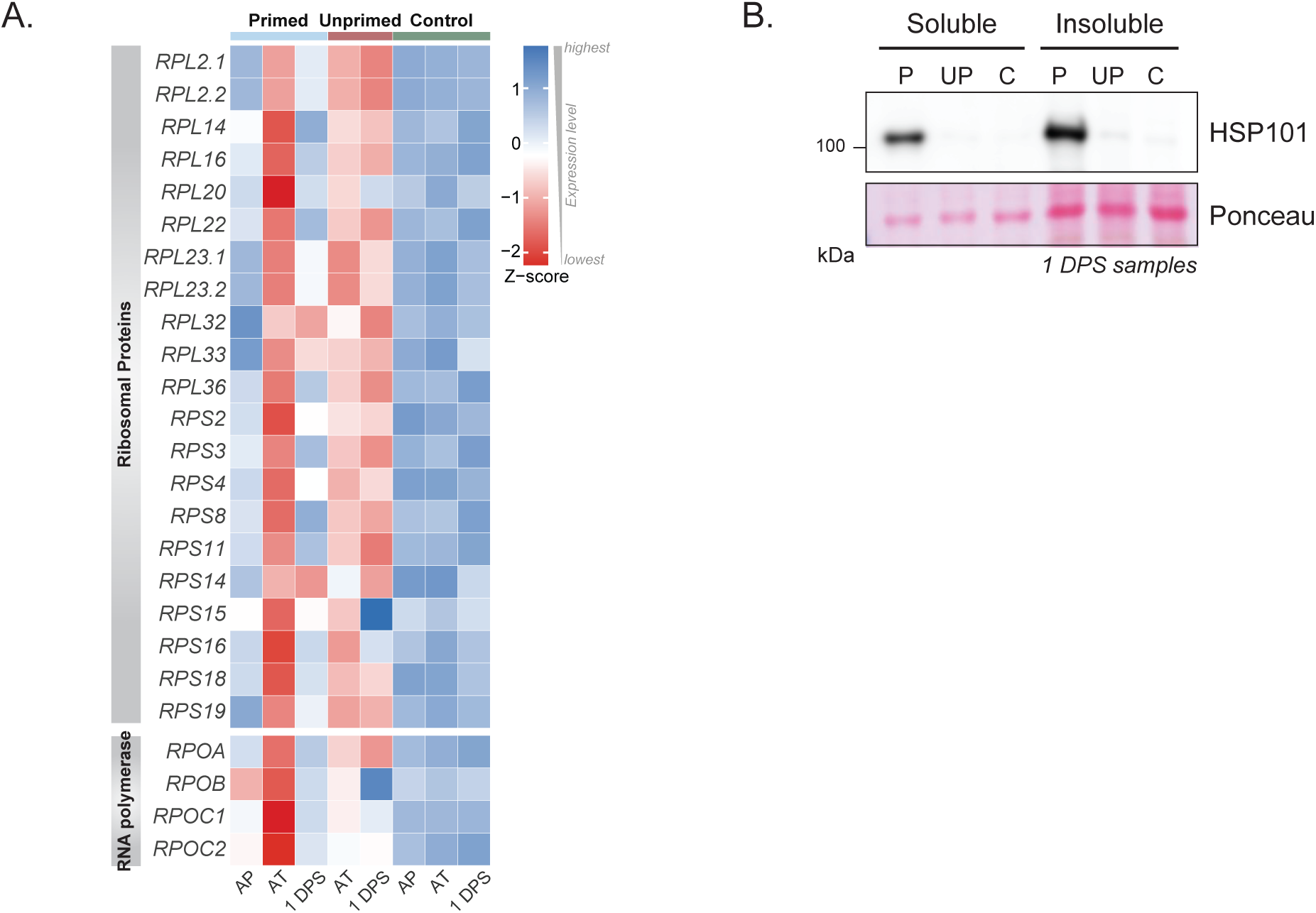
Unprimed seedlings experience translational arrest following triggering heat stress. (A) The heat-map shows the mean expression levels of differentially expressed genes associated with protein translation, including ribosomal proteins and RNA polymerases, derived from the transcriptome data. The priming stimulus alone (AP) does not alter the expression of these genes. However, upon triggering stress (AT), both primed and unprimed seedlings exhibited a marked downregulation, indicative of translational repression. During the recovery phase (1 DPS), primed seedlings restore the expression of translation-associated genes to near-control levels, whereas unprimed seedlings displayed reduced expression, suggesting a sustained translational arrest. (B) Immunoblot analysis showing induced levels of HSP101 in primed (both soluble and insoluble fractions) post-stress compared to control. HSP101 was undetectable in both the fractions of the unprimed seedlings, indicating an arrest of protein translation following heat stress (n=2 biological replicates). [AP - After priming; AT - After triggering (Day 0); DPS - Days post-stress, C - Control, P - Primed, UP - Unprimed].

**Figure S4:**
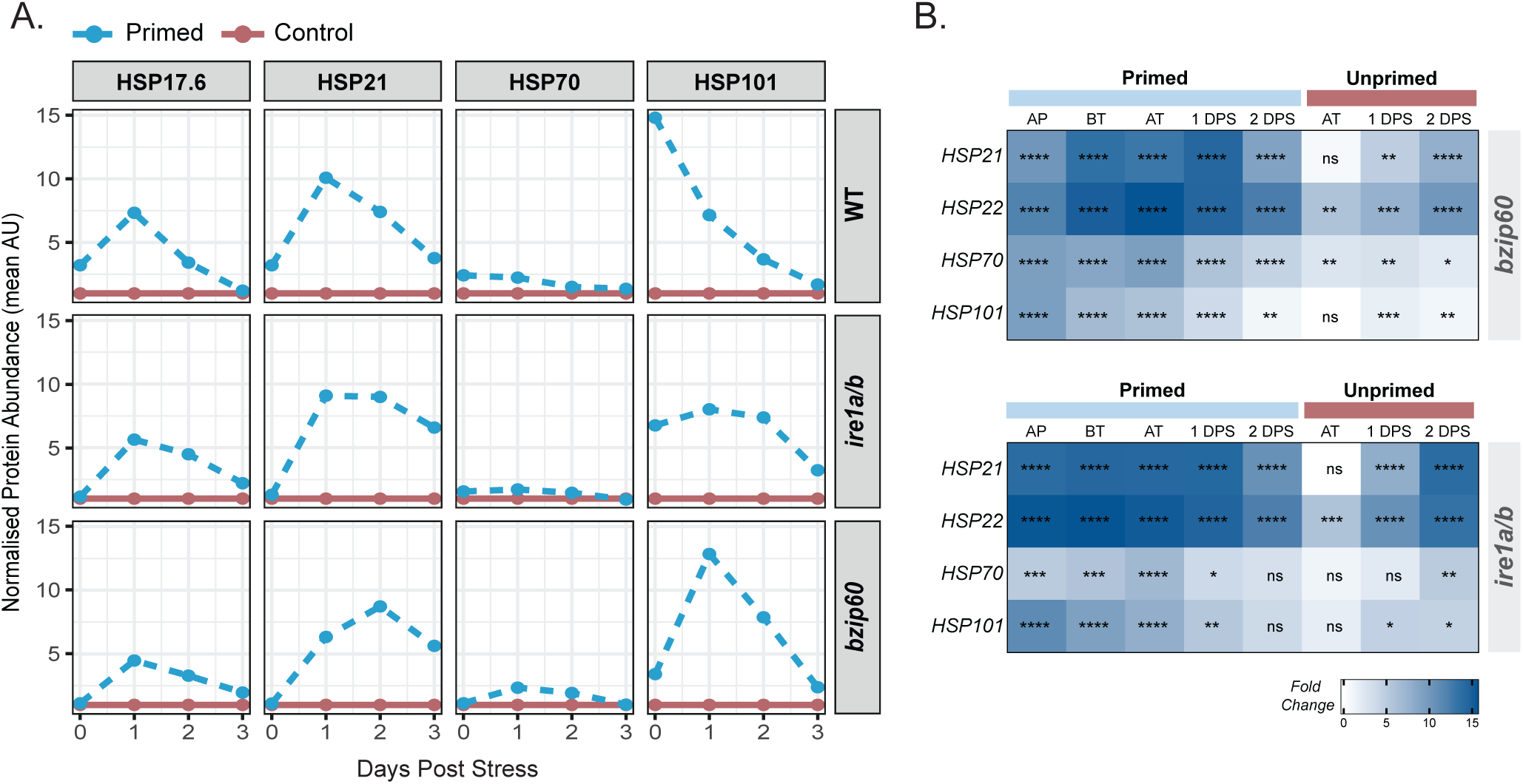
Loss of bZIP60 or IRE1A/B leads to delayed heat shock response. (A) Quantification of immunoblots revealed that HSP17.6 displays a similar pattern of increase in protein levels post-priming, across genotypes, compared to WT. A delay in the maximal induction of other HSPs was observed in the mutants compared to WT (n=3 biological replicates). (B) Heatmap represents the mean gene expression levels of *HSP21, HSP22, HSP70* and *HSP101* in *bzip60* and *ire1a/b* lines post-priming/triggering heat stress. In the primed, *HSPs* were highly induced upon priming and triggering, whereas, in the unprimed, induced expression of most of the *HSPs* was observed only by 1 DPS (n=2/3 biological replicates). Statistical significance was calculated using one-way ANOVA (**** p <0.0001; *** p <0.001; ** p <0.01; * p <0.05; ns – not significant) [AP - After priming, BT - Before triggering, AT - After triggering (Day 0), DPS - Days post-stress].

**Figure S5:**
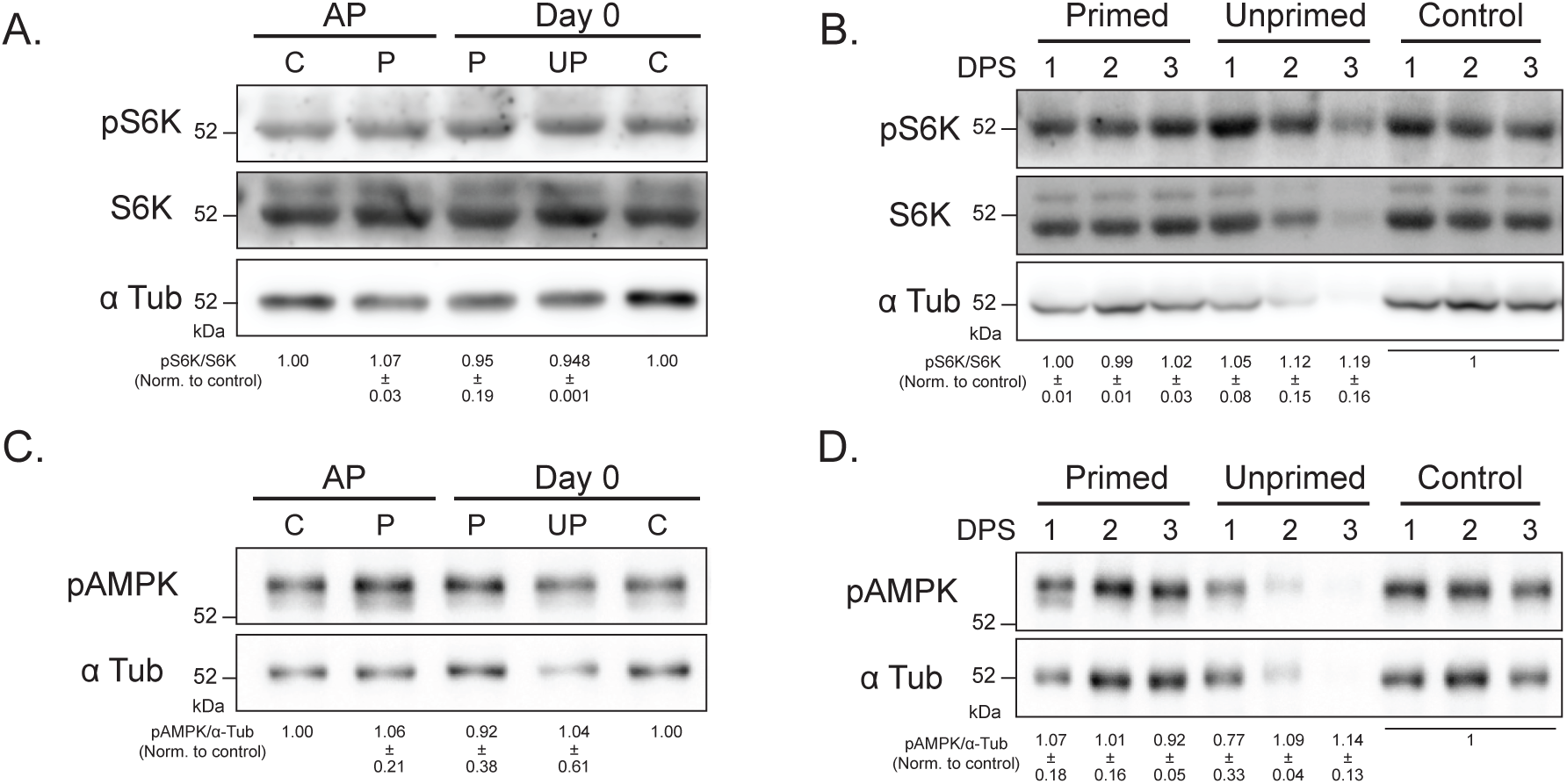
Priming-mediated autophagy response is TOR/SnRK independent. Immunoblot analysis of pS6K, S6K, and pAMPK under all the conditions at the given time points. (A-B) TOR kinase activity, as measured by pS6K/S6K levels, remained similar post-priming and triggering across conditions compared to control during priming/heat stress. The numbers below denote represents normalised mean intensity of pS6K/S6K respect to corresponding control time point (n=2 biological replicates). (C-D) The pAMPK/α-Tub levels do not change post-priming and triggering across conditions compared to control, indicating that SnRK activity remains similar during priming/heat stress. The numbers below denote represents normalised mean intensity of pAMPK/α-Tub respect to corresponding control time point (n=2 biological replicates). α-Tubulin served as a loading control. [AP - After Priming, DPS - Days Post Stress, C - Control, P - Primed, UP - Unprimed].

**Figure S6:**
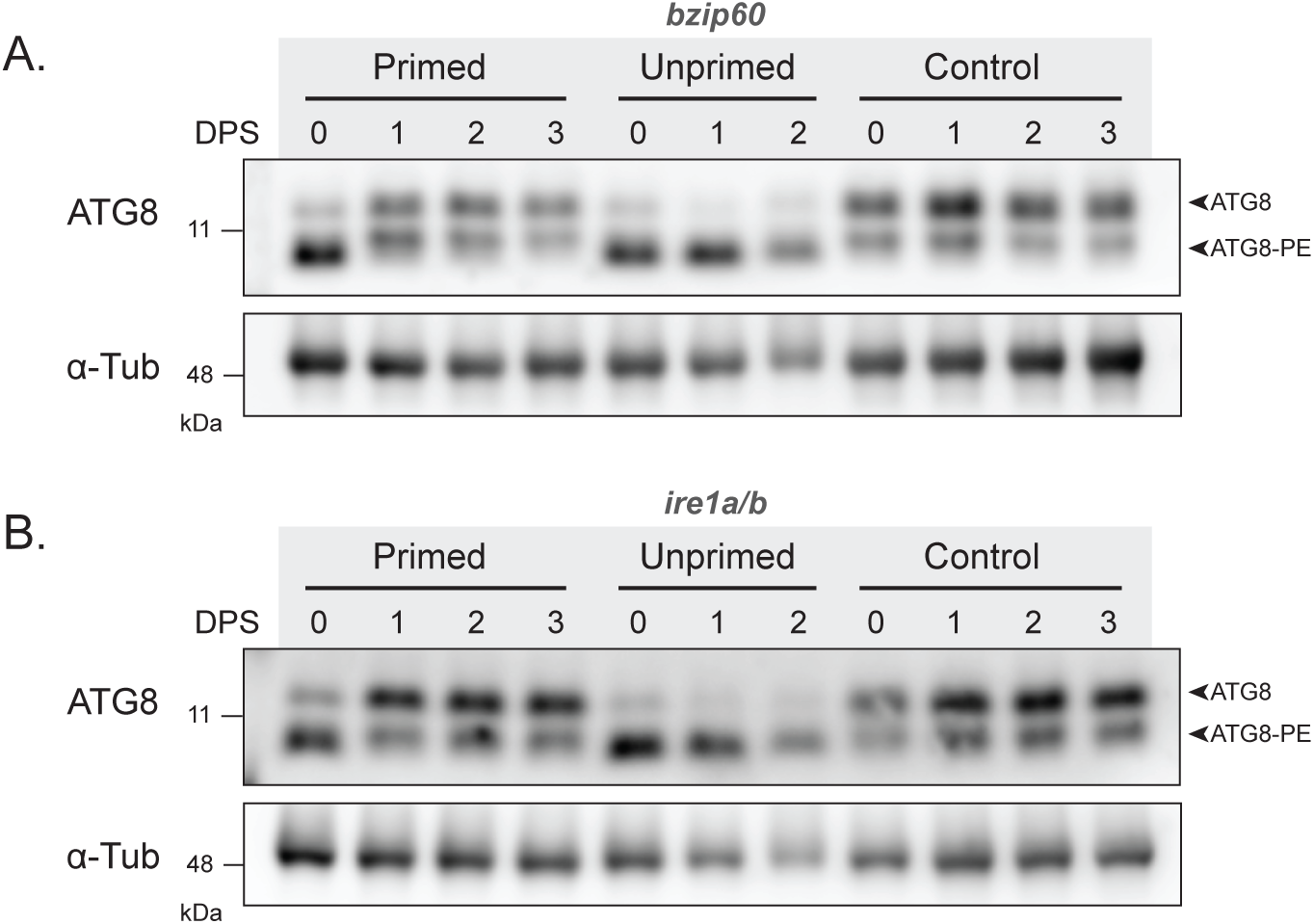
Autophagy remained induced in *bzip60* and *ire1a/b* mutants post-heat stress. Western blot analysis of ATG8 in (A) *bzip60* and (B) *ire1a/b* mutants post-heat stress revealed that autophagy was induced post-triggering (0 DPS) in primed seedlings; however, it reverted back to control levels during the stress recovery period (1-3 DPS). However, in unprimed (*bzip60* and *ire1a/b*), autophagy was induced post-triggering stress and stayed high even after removal of stress (n=3 biological replicates). α-Tubulin served as a loading control.

**Figure S7:**
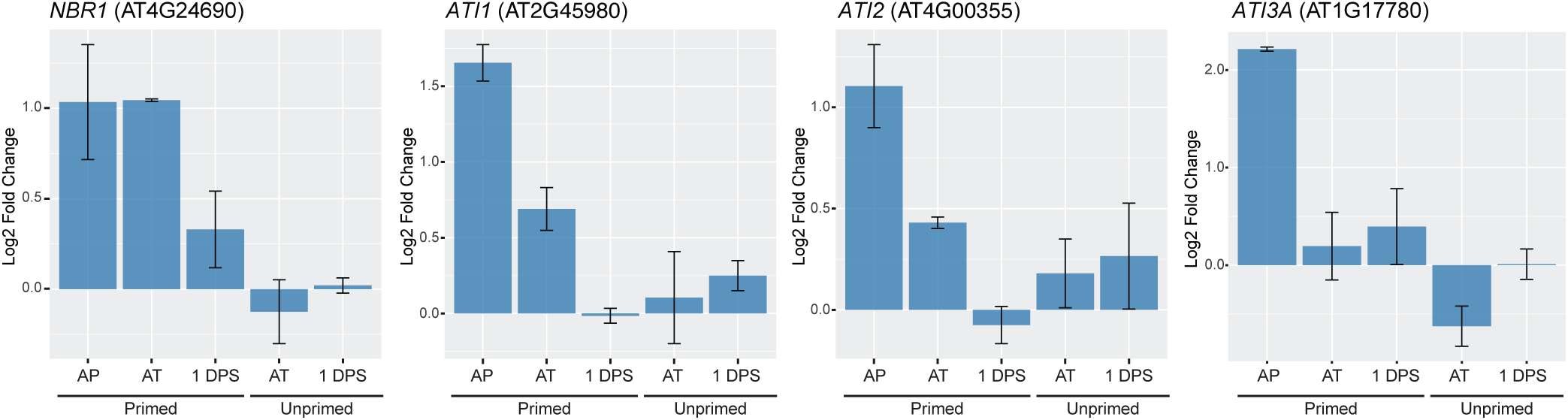
Transcript abundance of selective autophagy receptors in primed and unprimed. Relative transcript abundance of known selective autophagy receptors, *NBR1* (next-to-BRCA1) and ATG8-interacting protein 1 (*ATI1*), 2 (*ATI2*) and 3 (*ATI3*), was obtained from transcriptome data collected post-heat stress. All four genes were upregulated following priming stimulus (AP), whereas unprimed seedlings do not show change in expression. The preferential induction of these receptors suggests that selective autophagy may be activated in the primed seedlings following heat stress. [AP - After priming, AT - After triggering (Day 0), DPS - Days post-stress].

**Supplementary Table 1:**
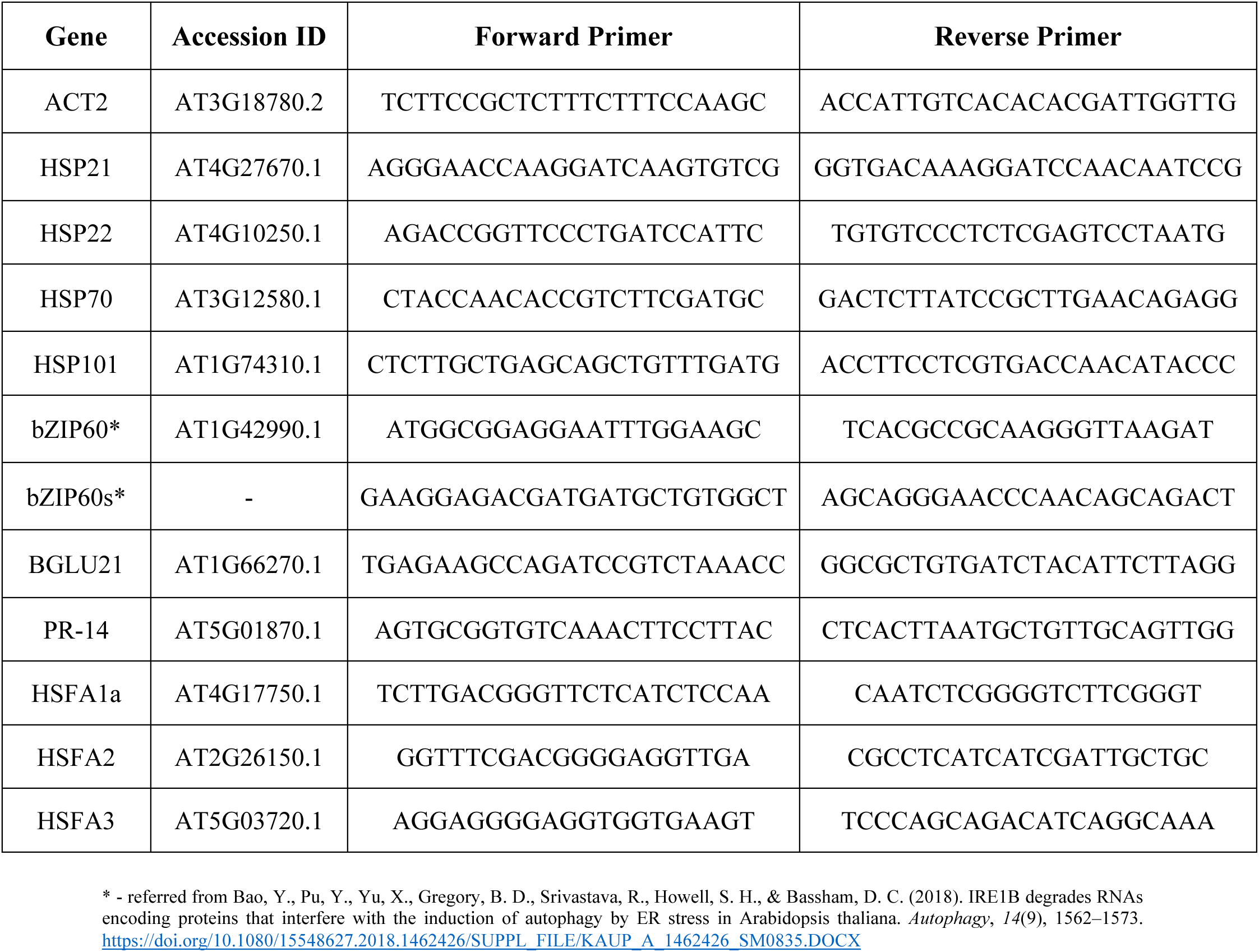
List of gene accession IDs and PCR primers.

